# Inhibition of S6K lowers age-related inflammation and immunosenescence and increases lifespan through the endolysosomal system

**DOI:** 10.1101/2022.08.25.505264

**Authors:** Pingze Zhang, James H. Catterson, Sebastian Grönke, Linda Partridge

## Abstract

The nutrient-sensing Target of Rapamycin complex 1 (TORC1) is an evolutionarily conserved regulator of longevity and healthspan. S6 kinase (S6K) is an essential downstream mediator for the effect of TORC1 on longevity. However, mechanistic insights on how TORC1-S6K signalling promotes lifespan and healthspan are still limited. Here we show that activity of S6K in the *Drosophila* fat body is essential for rapamycin-mediated longevity. Fat-body-specific activation of S6K blocked lifespan extension upon rapamycin feeding and induced accumulation of multilamellar lysosomal enlargements. By proteomics analysis we identified Syntaxin 13 (Syx13) as an important downstream mediator of TORC1-S6K signalling involved in regulating lysosomal morphology. Inhibition of TORC1-S6K signalling decreased age-associated hyperactivation of the NF-κB-like IMD pathway in the fat body and promoted bacterial clearance, mediated by Syx13, suggesting that lysosomal and immune function are connected during ageing. Middle-age-onset repression of IMD pathway in the fat body by Relish RNAi promoted bacteria clearance and extended fly lifespan. In mice, chronic rapamycin treatment elevated Syntaxin 12/13 (Stx12) level in liver. We identified alleviated immune processes in the aged liver as a common signature of S6K1-deficient and rapamycin-treated mice. Thus, our results indicate that suppression of the TORC1-S6K-Syx13 axis ameliorates both inflammageing and immunosenescence in hepatic tissues via the endolysosomal system and thereby extends longevity, providing a mechanistic explanation for the effects of rapamycin and suppression of S6K on immune function and lifespan in model organisms and, potentially, humans.

## Introduction

The mammalian Target of Rapamycin (mTOR) network senses nutrients and various stressors and plays a key role in the regulation of growth, metabolism and longevity. mTOR is a serine/threonine protein kinase which is a crucial component of mTOR Complex 1 (TORC1)^1^. Suppression of TORC1 signalling genetically or pharmacologically by feeding with FDA-licensed drug rapamycin, promotes lifespan and healthspan in various model organisms including nematode worms, fruit flies and mice^2-4^.

Rapamycin modulates lifespan via two evolutionary conserved TORC1 downstream effector mechanisms: autophagy and ribosomal S6 kinase (S6K)^3^. Autophagy is a lysosome-dependent cellular clearance pathway that is tightly linked with cellular and tissue homeostasis^5^. Promoting autophagy enhances longevity and suppresses age-related tissue deterioration in *Drosophila* and mice^5-7^. Moreover, recent findings implicate the intestine as a crucial organ mediating the effects of increased autophagy on health during ageing^*7-9*^. S6K is an AGC kinase family that regulates various fundamental cellular processes, including translation, lipid metabolism and immunity^10^. Mice deficient of S6K1 function are long-lived and resistant to age-related pathologies such as immune and motor dysfunction^11^. However, a mechanistic understanding of how TORC1-S6K signalling promotes lifespan and healthspan remains limited. *Drosophila melanogaster* provides an ideal model organism to dissect the role of S6K in regulating the ageing process as *Drosophila* has only a single S6K and it bears significant homology to the mammalian p70-S6K1^12^.

In addition to the lifespan extension, a further ageing-associated decline that is ameliorated by rapamycin is impaired immune function^13,14^. Age-associated changes in the immune system include a chronic inflammation, termed “inflammageing”, and deterioration of immune function, termed “immunosenescence”^15,16^. Age related changes in the immune system contribute to a vast range of age-associated diseases, such as cancer, diabetes, and cardiovascular diseases^16-18^. The nuclear factor-kappa B (NF-κB) pathway plays a key role in innate immunity and controls various aspects of immune responses. NF-κB signalling is tightly associated with the TORC1 pathway to modulate age-associated phenotypes. Rapamycin treatment ameliorates senescence-associated NF-κB activation^19^, and the effects of rapamycin in inflammageing and lifespan are diminished in mice with genetically enhanced NF-κB activity (*nf*κ*b1*^*−/−*^)^20^, suggesting the age-prolonging effects of rapamycin are partially mediated by its ability to limit NF-κB signalling. Moreover, late-life TORC1 inhibition using a rapamycin derivative improves immune function in older people after influenza vaccination, without significant adverse effects^14^. Interestingly, TORC1 pathway activity also shows significant associations with the load of bacteria in old flies^21,22^, suggesting that enhanced immune function from suppression of TORC1 is conserved in flies. However, the underlying molecular and cellular mechanisms of these TORC1-related effects are still elusive.

Here we uncovered mechanisms by which suppression of S6K activity ameliorates immune ageing and extends lifespan. We identified the *Drosophila* fat body as an essential organ to mediate S6K-dependent longevity. TORC1 inhibition by rapamycin treatment repressed enlarged multilamellar lysosomes in the *Drosophila* fat body, but this effect was blocked by elevating S6K activity. Importantly, we identified Syntaxin 13 (Syx13), a SNARE family protein that mediates endomembrane function, as a novel downstream mediator of TORC1-S6K signalling. Repressing Syx13 via RNAi mimicked S6K-dependent lysosomal enlargement, while over-expressing Syx13 rescued the S6K-dependent lysosomal enlargement and multilamellar structure in the fat body. We further identified that TORC1-S6K-Syx13 signalling regulated age-associated chronic inflammation and the decline in ability to clear bacteria via the endolysosomal pathway. Moreover, repression of inflammation in mid-adulthood, by knocking down *Drosophila* NF-κB like transcription factor Relish in the fat body, enhanced bacterial clearance and increased fly lifespan. In mice, rapamycin treatment increased expression of Syntaxin 12/13 (Stx12) in liver. Aggregating hepatic proteome and transcriptome of rapamycin-treated mice with the transcriptome of S6K1 deficient mice, we identified alleviated immune processes in the aged liver as common health outcome of TORC1-S6K inhibition. These findings define Syntaxin 12/13 as a novel downstream effector of TORC1-S6K signalling, highlighting a crucial role of the endolysosomal system in inflammageing, immunosenescence, and longevity.

## Results

### S6K activity in the fat body is essential for rapamycin-related longevity

To address whether downregulation of S6K activity prolongs lifespan in *Drosophila*, we used the inducible GeneSwitch system in combination with an S6K RNAi line to knock down S6K expression specifically in the adult stage. Ubiquitous downregulation of S6K activity using an *actGS* driver resulted in a small but significant increase in lifespan (Fig. 1a), demonstrating that, as in *C. elegans* and mice^11,23^, reduced S6K activity can also extend lifespan in *Drosophila*. To identify the tissue(s) in which reduced S6K activity acted to extend lifespan, we employed tissue-specific GeneSwitch drivers to suppress S6K activity in fat body tissue (*Lsp2GS*), neurons (*ElavGS*), muscle (*MHCGS*), heart tube (*HandGS*) and intestine (*TiGS*) of adult flies. Repression of S6K activity in neurons and intestine did not affect lifespan (Fig. S1a,d), while in muscle and heart tube it shortened it (Fig. S1b,c). Lifespan was extended only upon fat-body-specific repression of S6K activity (Fig. 1b) indicating that, in *Drosophila*, activity of S6K in the fat body is limiting for lifespan. As the lifespan extending effect of the TOR inhibitor rapamycin can be blocked by ubiquitous S6K activation^3^, we next tested whether activating S6K just in fat body tissue was sufficient to block the effect of rapamycin on survival. We administered rapamycin to flies that over-expressed a constitutively active S6K protein (*S6K*^*CA*^) under the control of the fat-body-specific *Lsp2GS* GeneSwitch driver and measured their survival. Similar to the ubiquitous activation (Fig. 1c), expression of *S6K*^*CA*^ specifically in the fat tissue was sufficient to block rapamycin-mediated lifespan extension (Fig. 1d). In summary, these results implicate the fat body as a key tissue regulating survival downstream of TORC1 and S6K.

**Figure 1:**
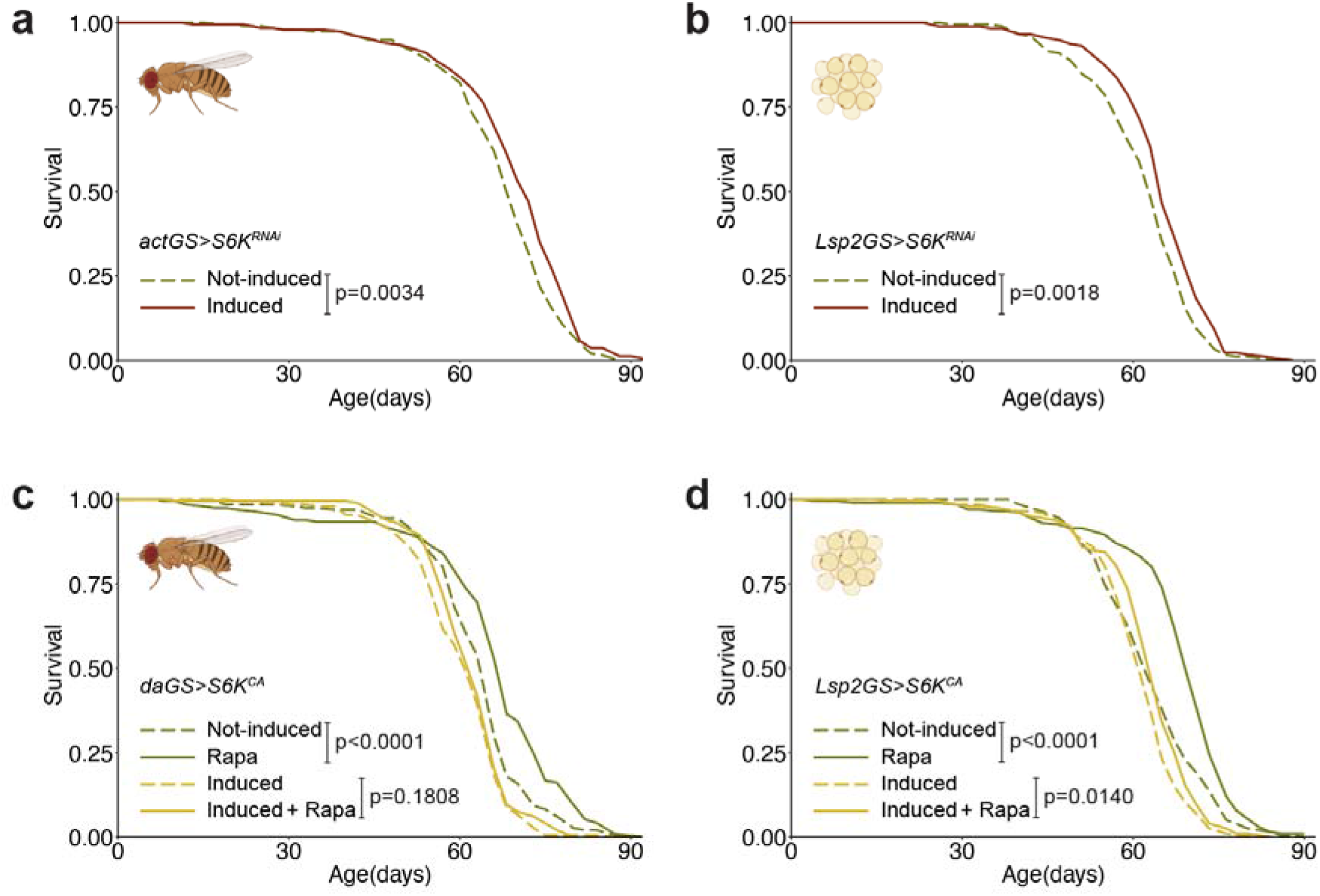
S6K activity in the fat body of adult flies determines longevity. **a**, Adult-onset repression of S6K ubiquitously using *actGS>S6K*^*RNAi*^ extended lifespan (n=200). **b**, Adult-onset repression of S6K in the fat body using *Lsp2GS>S6K*^*RNAi*^ extended lifespan (n=180). **c**, Rapamycin extended lifespan of control flies, but not of flies with ubiquitous overexpression of constitutively active S6K (*daGS>S6K*^*CA*^). Ubiquitous overexpression of constitutively active S6K significantly attenuated the response to rapamycin treatment (rapamycin: p<0.0001, *daGS>S6K*^*CA*^ induction: p<0.0001, interaction p=0.0236, n=200). **d**, Adult-onset S6K activation in the fat body (*Lsp2GS>S6K*^*CA*^) significantly attenuated rapamycin-related longevity (rapamycin: p<0.0001, *Lsp2GS>S6K*^*CA*^ induction: p<0.0001, interaction p=0.0331, n=200). Log-rank test and Cox Proportional Hazards (CPH) test.

### Proteomics analysis of fat body

The *Drosophila* fat body is functionally equivalent to mammalian liver and white adipose tissue and is the central organ for metabolism and immune responses^24^. To gain insight into the physiological actions of TORC1-S6K signalling in fat body during ageing, we used unbiased TMT-based proteomics profiling of fat bodies from day 10 (young) and day 50 (old) flies that over-expressed a constitutively active S6K protein (*S6K*^*CA*^) under the control of the fat body-specific *Lsp2GS* GeneSwitch driver (*Lsp2GS>S6K*^*CA*^) in combination with rapamycin treatment, or flies that over-expressed a S6K RNAi line to knock down S6K in the fat body (*Lsp2GS>S6K*^*RNAi*^) (Fig. S2a). Principal component analysis (PCA) revealed clear separation among age and treatment (Fig. S2b). We identified a total of 4101 proteins from the *Lsp2GS>S6K*^*CA*^ dataset and 4809 proteins from the *Lsp2GS>S6K*^*RNAi*^ dataset (Table 1). To explore S6K-dependent protein alterations, we filtered the proteins that were significantly changed (p<0.05) by both rapamycin treatment and S6K manipulation. Proteins regulated by TORC1-S6K signalling were associated with endocytosis, RNA binding processes and extracellular matrix in young fat bodies, while in old fat bodies mitochondria-, translation-and immune-related proteins were significantly altered (Fig. S2c, Table 2).

To further enhance the resolution of our analysis, we conducted network propagation analysis^25^ to incorporate information on protein-protein interactions. We then identified functional categories and clustered them into rapamycin-induced, S6K-dependent terms and S6K inhibition-induced terms (Fig. S2d). Proteins directly affected by TORC1-S6K signalling were endosome/lysosome-oogenesis-, and translation-related, while at later age immune-, translation-and lipid-related processes were altered.

### S6K activity in the fat body does not affect fecundity, global translation or triacylglycerol homeostasis

The fat body plays an important role in oogenesis and reproduction in flies and reduced reproduction is often associated with increased longevity^26^. As oogenesis was identified as a significantly enriched term in the proteomics analysis, we measured egg laying of females with fat body specific S6K downregulation and found that fecundity was not affected (Fig. S3a). In addition, repression of S6K activity significantly extended survival of sterile females carrying the *ovo*^*D*^ mutation (Fig. S3b). These results suggest, that reduced reproduction is not a causal factor in S6K-related lifespan-extension, consistent with the findings for lifespan extension by rapamycin^3^.

S6K is a regulator of protein synthesis, and protein translation was identified as an enriched term in the proteomics analysis. Furthermore, reduced protein synthesis has been associated with longevity in several invertebrate longevity models. Thus, we addressed whether S6K or rapamycin affect global protein synthesis in the fly fat body by performing puromycin incorporation assays. Interestingly, puromycin incorporation was affected neither by rapamycin treatment nor by S6K activation upon rapamycin feeding (Fig. S4), suggesting that chronic rapamycin treatment and S6K activity do not change global translation rates in the fly fat body. Consistently, chronic rapamycin feeding had also no effect on global translation rates in multiple mouse tissues^27^.

Lipid metabolism has been implicated in the ageing process^28^ and rapamycin has been shown to increase lipid storage in flies^3^. Therefore, we investigated whether S6K was involved in regulation of lipid metabolism. However, S6K activation did not alter rapamycin-induced triglyceride accumulation in the fat body (Fig. S5a). In addition, repressing S6K in the fat body did not affect survival under starvation (Fig. S5b), and ubiquitous S6K activation did not attenuate rapamycin-induced starvation resistance (Fig. S5c). Thus, TORC1-S6K-dependent longevity cannot be explained by the regulation of lipid homeostasis.

### TORC1-S6K signalling in the fat body affects lysosomal morphology

Proteomics profiling showed that lysosome-related annotations were associated with TORC1-S6K activity in the young fat body (Fig. S2d). To explore the role of TORC1-S6K signalling on lysosomes, we stained fat bodies with lysotracker, a fluorescent, acidotropic dye that stains lysosomes and autophagy-associated autolysosomes^29^. In line with the role of TORC1 in lysosomal biogenesis and autophagy, rapamycin treatment increased the number of lysotracker-positive puncta in fat body cells. Interestingly, S6K activation did not attenuate rapamycin-induced, lysotracker-positive puncta but led to a substantial accumulation of enlarged acidic organelles in fat body cells (Fig. 2a). These S6K-dependent enlarged organelles were further identified as lysosomes, indicated by a *GFP-Lamp1* reporter^30^, and late endosomes, indicated by a *YFP-Rab7* reporter (Fig. 2b). To measure the change in lysosomes directly, we used a *Lamp1-3xmCherry* reporter^31^, which marks lysosomes by an endogenous promoter-driven C-terminally mCherry-tagged Lamp1. Consistent with the results from lysotracker staining, mCherry-labelled lysosomes were enriched as large puncta in the fat bodies with S6K activation co-treated with rapamycin (Fig. S6a). In addition, these enlarged lysosomes presented as multilamellar structures when analysed by electron microscopy (Fig. 2c). Of note, the ratio of multilamellar lysosomes was ameliorated by rapamycin treatment in an S6K-dependent manner. Thus, TORC1-S6K signalling regulated changes in lysosomal morphology in the fat body.

**Figure 2:**
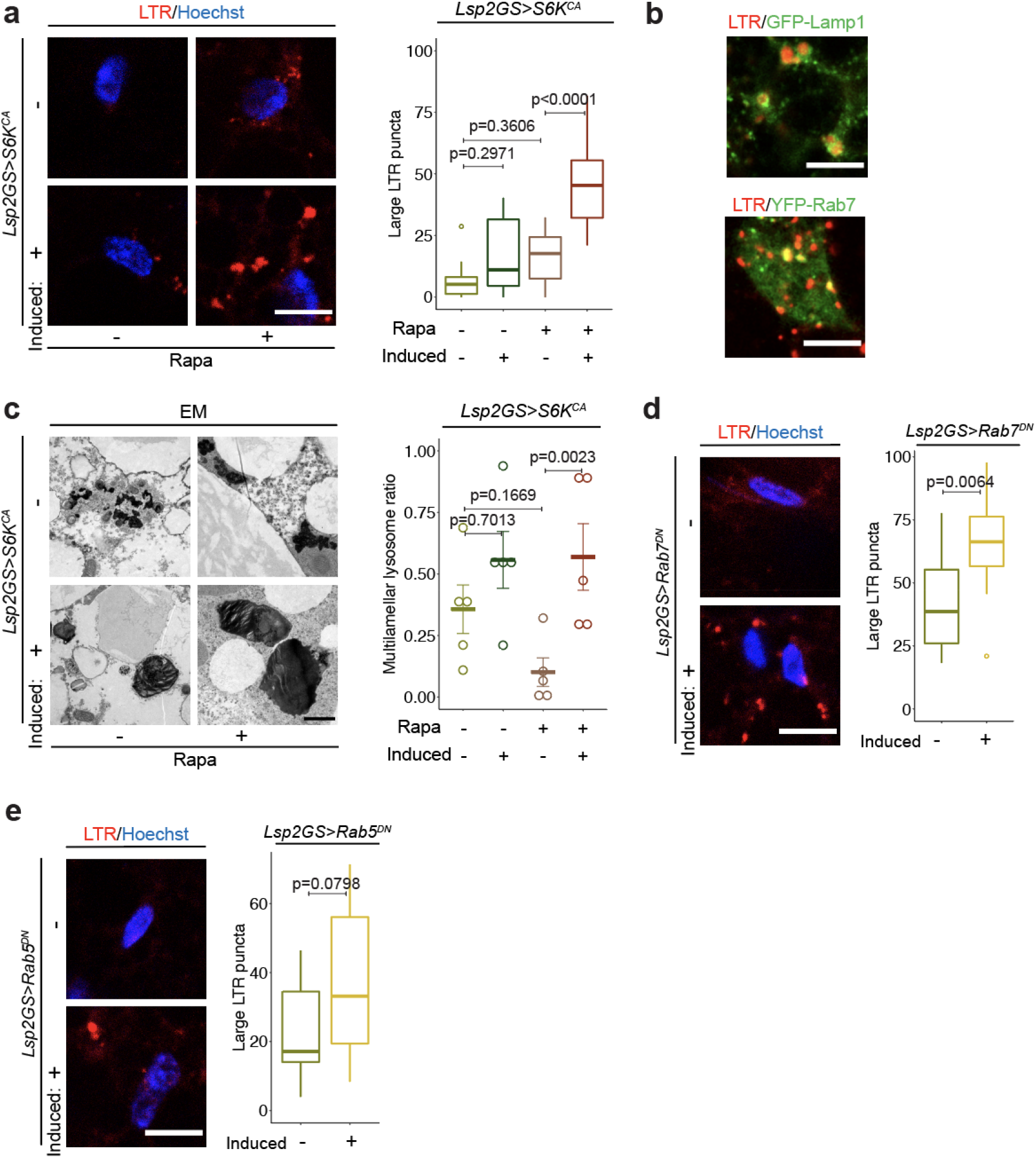
TORC1-S6K signalling affects lysosomal morphology in the fat body. **a**, Lysotracker staining of fat bodies from young (day 10) flies treated with rapamycin and overexpressing constitutively active S6K (*Lsp2GS>S6K*^*CA*^). Fat body-specific adult-onset overexpression of constitutively active S6K significantly increased acidic organelle size in response to rapamycin treatment (rapamycin: p<0.0001, *Lsp2GS>S6K*^*CA*^ induction: p<0.0001, interaction p=0.0246, n=12). **b**, Lysotracker-positive enlarged acidic organelles (red) colocalized with the lysosomal marker Lamp1 (*Lsp2GS>S6K*^*CA*^;*GFP-Lamp1*, green, upper panel) and partially colocalized with Rab7 (*Lsp2GS>S6K*^*CA*^;*YFP-Rab7*, green, lower panel), a marker for late endosomes, in young flies treated with rapamycin and overexpressing S6K^CA^. **c**, Electron microscopy imaging of fat bodies of young flies treated with rapamycin and overexpressing S6K^CA^. Overexpression of S6K^CA^ in the fat body attenuated the effect of rapamycin on multilamellar lysosomes (rapamycin: p=0.0395, *Lsp2GS>S6K*^*CA*^ induction: p=0.2798, interaction p=0.0388, n=5). **d**, Lysotracker staining of young fat bodies overexpressing dominant negative Rab7 (*Lsp2GS>Rab7*^*DN*^). Fat body-specific overexpression of Rab7^DN^ significantly increased acidic organelle size (p=0.0064, n=12). **e**, Fat body-specific overexpression of dominant negative Rab5 (*Lsp2GS>Rab5*^*DN*^) also increased acidic organelle size (p=0.0798, n=12). Data are displayed as Tukey box plot **(a, d-e)** or mean□±□s.e.m. **(c**). Each data point represents an average value per fat body. Scale bar, 10 μm **(a-b, d-e)** or 1 μm **(c)**. Linear mixed model **(a, d-e)** or negative binomial generalized linear model **(c)** followed by Tukey’s multiple comparison test.

The lysosome is the terminal compartment for autophagy and endocytosis^32^. As autophagy plays a vital role during ageing^5^, we firstly investigated if a defective autophagy process caused the TORC1-S6K-dependent lysosomal changes. Consistent with previous findings^3^, repressing autophagy induction by *Lsp2GS>Atg5*^*RNAi*^ blocked rapamycin-induced accumulation of lysotracker-positive organelles in the fat body (Fig. S7a), indicative of perturbed autophagy. Surprisingly, the number of enlarged lysotracker-positive organelles induced by S6K activation with rapamycin treatment was not changed upon inhibition of Atg5 in the fat body (Fig. S7b), suggesting that the S6K-dependent lysosomal enlargement was not caused by autophagy dysfunction.

TORC1-S6K-dependent enlarged lysotracker-positive organelles were partially colocalised with late endosomes (Fig. 2b). Thus, we next investigated if the endocytosis pathway might contribute to the S6K-dependent lysosomal enlargement. We impaired late endosome formation by expressing dominant-negative (DN) Rab7 (*Lsp2GS>Rab7*^*DN*^), which led to enlarged lysotracker-positive organelles (Fig. 2d). Moreover, perturbation of early endosomes by Rab5 DN (*Lsp2GS>Rab5*^*DN*^) also mildly induced lysotracker-positive organelle enlargement (Fig. 2e). These results suggest that the endocytosis pathway could contribute to TORC1-S6K-related lysosomal morphological changes in the fat body.

### Syntaxin 13 is a downstream effector of TORC1-S6K signalling regulating lysosomal structure

The presence of multilamellar lysosomes suggested impaired lysosomal fusion or defective membrane function, which is typical of lysosomal storage diseases^33^. As vesicle fusion and vesicle membrane-related processes were negatively regulated by TORC1-S6K signalling (Fig. S2d), Syntaxin 13 (Syx13), the most differentially abundant S6K-dependent SNARE protein, was identified as a promising TORC1-S6K downstream effector (Fig. 3a-b, Table 2). RNAi-mediated repression of Syx13 in young fat bodies recapitulated S6K-associated lysosomal enlargement (Fig. 3c). However, overexpressing Syx13 did not alter lysosomal morphology under basal condition (Fig. 3d). To determine whether Syx13 was causally associated with the S6K-induced changes in lysosomal morphology, we increased both Syx13 expression and S6K activity in the fat body of flies that were treated with rapamycin. Strikingly, S6K-induced enlarged lysosomes (Fig. 3e) and the associated multilamellar structure (Fig. 3f) were diminished by Syx13 over-expression. Therefore, TORC1-S6K-Syx13 signalling mediated the lysosomal structural changes.

**Figure 3:**
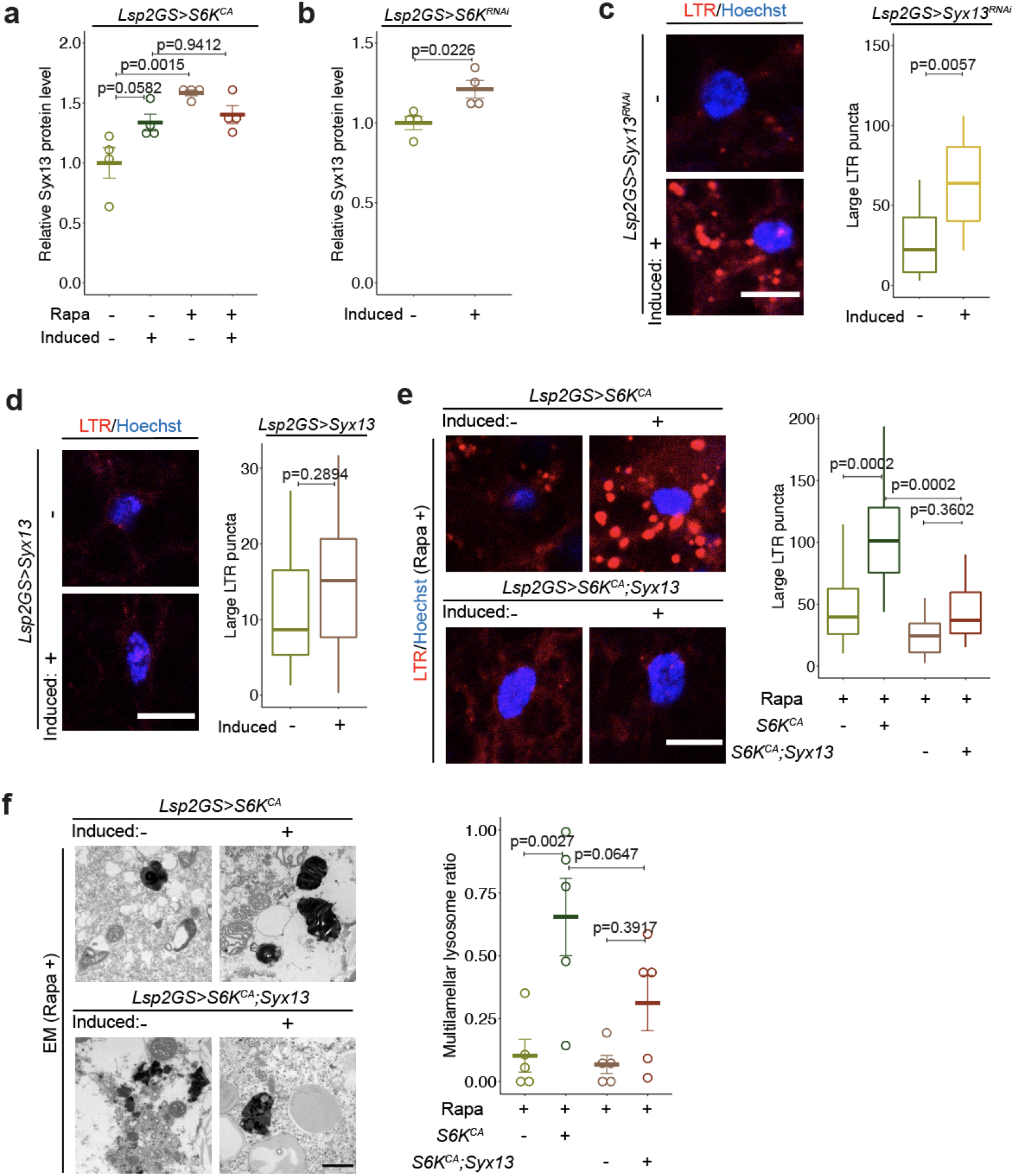
Syntaxin 13 is a downstream effector of TORC1-S6K signalling that regulates lysosomal structure in the fly fat body. **a**, Syx13 protein level was increased upon rapamycin treatment in young fat body cells and this increase was partly reverted by S6K overexpression (*Lsp2GS>S6K*^*CA*^; rapamycin: p=0.0003, *Lsp2GS>S6K*^*CA*^ induction: p=0.0138, interaction p=0.0086, n=5). **b**, Syx13 protein level was increased upon S6K repression (*Lsp2GS>S6K*^*RNAi*^; p=0.0226, n=5) in young fat body cells. Syx13 protein levels were measured by mass spectrometry-based proteomics. **c**, Knock-down of Syx13 expression (*Lsp2GS>Syx13*^*RNAi*^) resulted in enlarged lysosomes in the fat body of young flies, depicted by lysotracker staining (p=0.0057, n=14). **d**, Overexpression of Syx13 (*Lsp2GS>Syx13*) did not affect lysosomal enlargement (p=0.2894, n=12) in young fat bodies. **e**, Overexpression of Syx13 (*Lsp2GS>S6K*^*CA*^*;Syx13*) rescued the enlarged lysosomes of flies overexpressing S6K (*Lsp2GS>S6K*^*CA*^) treated with rapamycin (*Lsp2GS>S6K*^*CA*^ induction: p<0.0001, *Lsp2GS>Syx13* induction: p<0.0001, interaction p=0.0429, n=12). **f**, Overexpression of Syx13 (*Lsp2GS>S6K*^*CA*^*;Syx13*) partially rescued the multilamellar lysosomes of S6K overexpressing (*Lsp2GS>S6K*^*CA*^) flies treated with rapamycin, depicted by electron microscopy (*Lsp2GS>S6K*^*CA*^ induction: p=0.0005, *Lsp2GS>Syx13* induction: p=0.7989, interaction p=0.3068, n=5). Data are displayed as mean□±□s.e.m. **(a, b, f)** or displayed as Tukey box plot **(c-e)**. Each data point represents an average value per five fat bodies **(a-b**) or per fat body **(c-f)**. Scale bar, 10 μm **(c-e)** or 1 μm **(f)**. Linear mixed model **(a-e)** or negative binomial generalized linear model **(f)** followed by Tukey’s multiple comparison test.

### Inflammageing and Immunosenescence are modulated by TORC1-S6K signalling

A growing body of evidence has established that the ageing immune system can lead to a low-grade inflammation which is associated with increased mortality^18,34,35^. The fat body is a major immune organ of *Drosophila*, and the proteins in old fat bodies that were altered by TORC1-S6K signalling showed significant enrichment of immune-related GO annotations (Fig. S2d). The primary immune response pathway in *Drosophila* is immune deficiency (IMD) signalling. Upon activation of the IMD pathway, the NF-κB-like transcription factor Relish is cleaved in the cytoplasm, and the activated part of Relish translocates to the nucleus and induces the transcription of antimicrobial peptides (AMPs), such as *Diptericin A (DptA)* ^36-38^. In line with previous reports^37,39,40^, aged fat bodies exhibited elevated Relish cleavage (Fig. 4a), Relish-positive nuclei (Fig. 4b) and *DptA* expression (Fig. 4c), indicating that IMD signalling was activated during ageing. These changes were significantly ameliorated by reduced S6K activity in the fat body (Fig. 4d-f). Furthermore, rapamycin treatment greatly suppressed the age-associated changes in Relish localisation and *DptA* expression in the fat body, while activation of S6K blocked rapamycin-related changes in IMD signalling (Fig. 4g-h). Another common feature of the aged immune system is immunosenescence, represented by a declining ability of old flies to clear pathogens^41,42^. We therefore investigated whether S6K plays a role in bacterial clearance upon systemic infection with *Ecc15*, a gram-negative bacterium widely used to study *Drosophila* immune responses^43^. Rapamycin treatment increased bacterial clearance in old flies, and expression of *S6K*^*CA*^ specifically in the fat tissue blocked this effect (Fig. 4i). These results suggest that TORC1-S6K signalling regulates age-associated activation of the IMD pathway and the decline of pathogen clearance.

**Figure 4:**
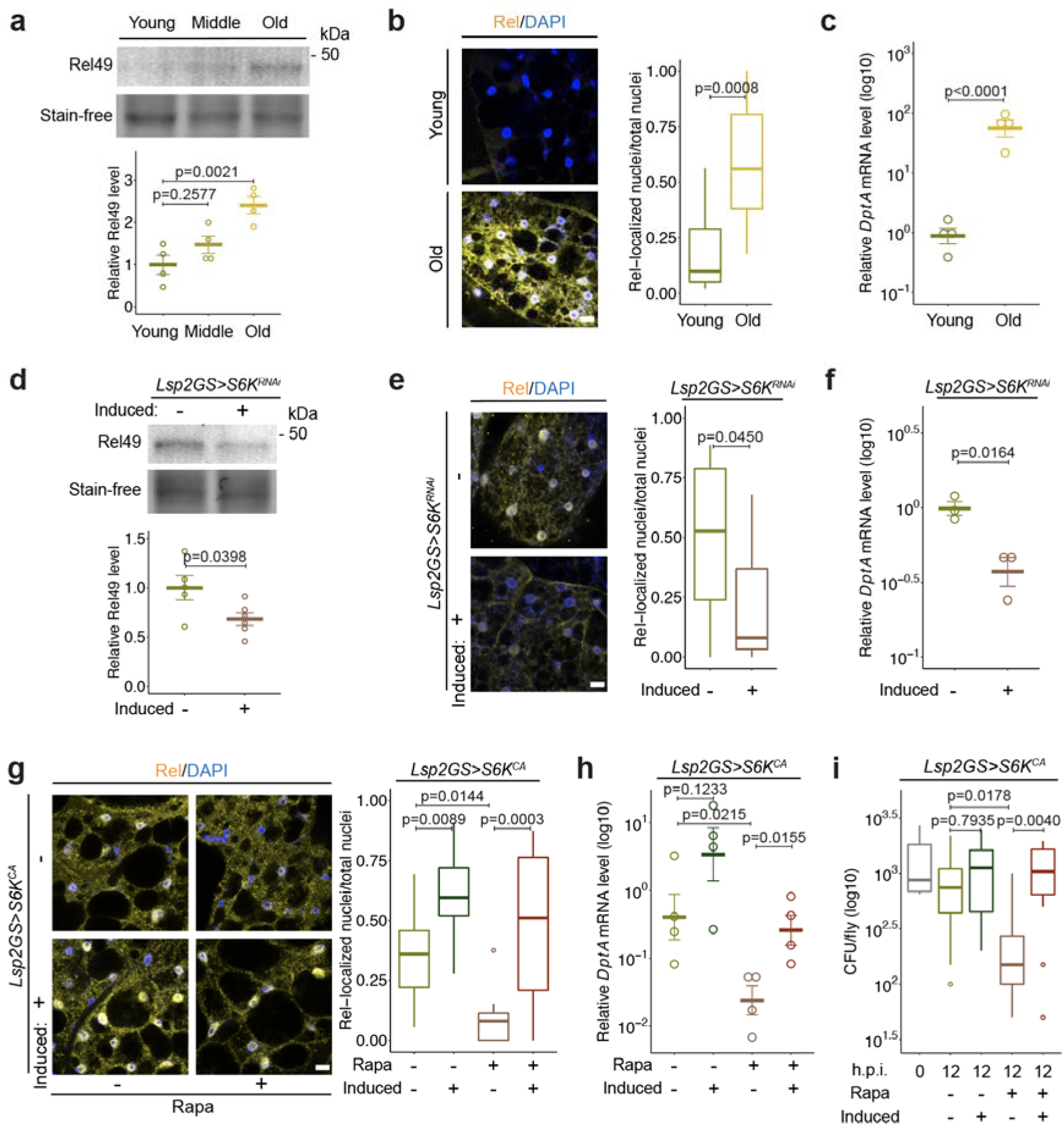
Immune ageing is modulated by TORC1-S6K signalling in the fat body. **a**, Cleaved Relish (Rel49, 49 kDa) in fat bodies of young (day 10), middle (day 30), and old (day 50) flies (age effect p=0.0034, n=4). **b-c**, Relish protein localisation (**b**, n=9 in young and n=13 in old) and *DptA* transcript expression (**c**, n=4) in fat bodies of young and old flies. **d-f**, S6K inhibition (*Lsp2GS>S6K*^*RNAi*^) suppressed the age-related increase in activated Rel49 (**d**, n=5), accumulation of Relish in the nucleus (**e**, n=14), and the increase in *DptA* expression (**f**, n=3). **g-h**, Rapamycin treatment suppressed age-related Relish localisation **(g)** and the increase in *DptA* **(h)**. Overexpression of S6K (*Lsp2GS>S6K*^*CA*^) blocked the effect of rapamycin on Relish localisation (**g**, rapamycin: p=0.0025, *Lsp2GS>S6K*^*CA*^ induction: p<0.0001, interaction p=0.2345, n=14) and *DptA* expression (**h**, n=4). **i**, Rapamycin treatment improved bacterial clearance in old flies infected with *Ecc15*. This effect was blocked by S6K overexpression (*Lsp2GS>S6K*^*CA*^) (rapamycin: p=0.0185, *Lsp2GS>S6K*^*CA*^ induction: p=0.0024, interaction p=0.0620, n=12). Data are displayed as mean□±□s.e.m. **(a, c-d, f, h)** or Tukey box plot **(b, e, g, i)**. Each data point represents an average value per fat body **(b, e, g**), per five fat bodies **(a, c, d, f, h)**, or per three whole flies **(i)**. Scale bar, 10 μm. One-way ANOVA followed by Dunnett’s multiple comparison test **(a);** linear mixed model **(b, e, g)** or two-way ANOVA with log transformation **(i)** followed by Tukey’s multiple comparison test; two-sided Student’s t-test with **(c, f, h)** or without log-transformation **(d)**.

Potential mechanisms of age-associated immune activation include increased intestinal permeability, chronic infection, and “sterile inflammation” caused by internal factors such as cellular senescence and oxidative stress ^18,44^. We next investigated whether S6K directly regulates IMD activation upon microbial infection. We therefore challenged young fat bodies *ex vivo* with *Ecc15*. Although *Ecc15* infection led to a clear induction in the number of Relish-positive nuclei under basal conditions, knock-down of S6K failed to repress this induction in the young fat body (Fig. S8a), suggesting that the effect of S6K on the IMD pathway is indirect and dependent on age-dependent changes in the fly fat body. We next explored whether activation of the IMD pathway by TORC1-S6K signalling is dependent on age-associated accumulation of bacteria^45^. Therefore, flies were treated with an antibiotic cocktail from young age to eliminate bacterial accumulation during ageing. Colony unit forming assays indicated that the antibiotic cocktail dramatically reduced the total number of microbes present (Fig. S8b). However, rapamycin treatment still repressed the number of Relish-positive nuclei and *DptA* expression in the fat body in an S6K-dependent manner (Fig. S8c-d). Taken together, these results show that TORC1-S6K signalling regulates age-associated activation of the IMD pathway in the fat body independent of bacterial load.

### The endolysosomal system mediates the effects of rapamycin on inflammageing and immunosenescence

Lysosomal function plays a crucial role in inflammatory and autoimmune disorders^46^. Given that TORC1-S6K-Syx13 signalling regulated lysosomal morphology, we hypothesised that age-related IMD activation in the fat body could be an outcome of endolysosomal dysfunction. We therefore first investigated if the endocytosis pathway contributes to TORC1-S6K-related regulation of Relish. Interestingly, over-expression of dominant-negative (DN) Rab7 blocked the rapamycin-induced decrease in the number of Relish-positive nuclei in the aged fat body (Fig. 5a). Moreover, impairment of early endosomes by expression of Rab5 DN also attenuated TORC1-related Relish repression (Fig. 5b), suggesting that the endolysosomal system acts downstream of TORC1-S6K in regulating IMD activity.

**Figure 5:**
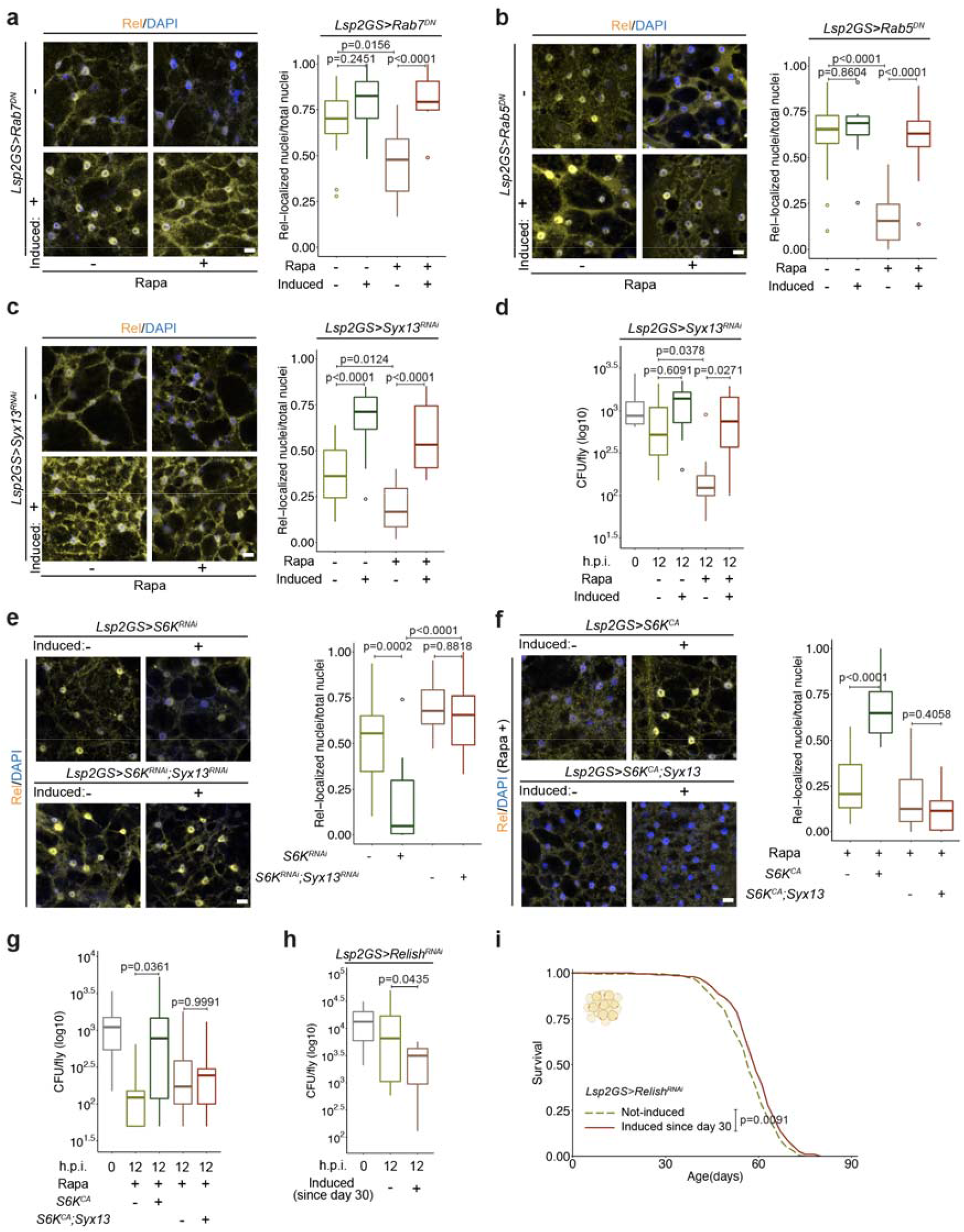
The TORC1-S6K-Syx13 axis regulates immunoageing via the endolysosomal system. **a-c**, Rapamycin treatment suppressed the age-related nuclear localisation of Relish in old fat body cells. Dominant negative Rab7 (**a**, rapamycin: p=0.0638, *Lsp2GS>Rab7*^*DN*^ induction: p<0.0001, interaction p=0.0160, n=14) and Rab5 (**b**, rapamycin: p<0.0001, *Lsp2GS>Rab5*^*DN*^ induction: p<0.0001, interaction: p=0.0003, n=14) blocked the effect of rapamycin on the age-related nuclear localisation of Relish. **c-d**, Knockdown of Syx13 (*Lsp2GS>Syx13*^*RNAi*^) blocked the effect of rapamycin on (**c**) Relish nuclear localisation (rapamycin: p=0.0012, *Lsp2GS>Syx13*^*RNAi*^ induction: p<0.0001, interaction p=0.2930, n=14) and (**d)** bacterial clearance (rapamycin: p=0.0093, *Lsp2GS>Syx13*^*RNAi*^ induction: p=0.0057, interaction p=0.2226, n=8). **e**, Knockdown of Syx13 (*Lsp2GS>S6K*^*RNAi*^*;Syx13*^*RNAi*^) blocked the effect of S6K knockdown on Relish nuclear localisation (*Lsp2GS>S6K*^*RNAi*^ induction: p=0.0004, *Lsp2GS>Syx13*^*RNAi*^ induction: p<0.0001, interaction: p=0.0088, n=14). **f-g**, Overexpression of Syx13 (*Lsp2GS>S6K*^*CA*^*;Syx13*) rescued the effect of S6K activation on Relish localisation (**f**, *Lsp2GS>S6K*^*CA*^ induction: p<0.0001, *Lsp2GS>Syx13* induction: p<0.0001, interaction p<0.0001, n=14) and bacterial clearance (**g**, *Lsp2GS>S6K*^*CA*^ induction: p=0.0644, *Lsp2GS>Syx13* induction: p=0.7042, interaction: p=0.0429, n=10) in old flies treated with rapamycin. **h-i**, Middle-age-onset repression of Relish using *Lsp2GS>Relish*^*RNAi*^ improved bacterial clearance (**h**, n=8 in 0 h.p.i. group and n=12 in 12 h.p.i. groups) and extended lifespan (**i**, p=0.0091, n=200). Data are displayed as Tukey box plot. Each data point represents an average value per fat body **(a-c, e-f)** or per three whole flies **(d, g-h)**. Scale bar, 10 μm. Linear mixed model **(a-c, e-f)**, two-way ANOVA with log transformation **(d, g)** followed by Tukey’s multiple comparison test; two-sided Student’s t-test **(h)**; log-rank test **(i)**.

We next evaluated if Syx13 acted downstream of TORC1-S6K to regulate immunosenescence. Knock down of Syx13 caused a robust induction in the number of Relish-positive nuclei in the old fat body and blocked the effects of rapamycin on Relish-positive nuclei and bacterial clearance (Fig. 5c-d). Relish repression upon S6K knock down was also rescued by reducing Syx13 expression (Fig. 5e). Furthermore, while S6K activation blocked the effects of rapamycin on Relish-positive nuclei and bacterial clearance in the old fat body, co-over-expression of Syx13 restored the effects of rapamycin (Fig. 5f-g). Thus, Syx13 acts downstream of TORC1 and S6K to regulate immunosenescence in the ageing fat body. Despite the finding that autophagy can control inflammation and longevity^47^, blocking autophagy by Atg5 RNAi in the fat body did not block rapamycin-dependent repression of Relish-positive nuclei and longevity (Fig. S7c-d). Taken together, these findings show that TORC1-S6K-Syx13 signalling regulates immune ageing via the endolysosomal system.

### Reduced relish activity in the fat body prevents immunosenescence and increases longevity

Given that TORC1-S6K signalling regulated age-associated immune dysfunctions, we next addressed whether age-associated inflammation in the fly fat body affects pathogen clearance and longevity. Down-regulating Relish level via RNAi in the fat body from middle-age on significantly improved bacterial clearance and extended fly lifespan (Fig. 5h-i), demonstrating that ameliorating inflammageing may contributes to efficient clearance of bacteria at old age and lifespan extension.

### TORC1-S6K signalling regulates Syntaxin 12/13 expression and immunoageing in mouse liver

Syx13 levels were upregulated in the fly fat body upon downregulation of TORC1-S6K signalling. Interestingly, Syx13 levels were also upregulated in long-lived S6K mutant worms^23^ and in rapamycin-treated yeast^48^, suggesting that this regulation is evolutionarily conserved, at least among yeast, worms and flies. To assess if the regulation of Syx13 by rapamycin is also conserved in mammals, we used western blot to measure Syntaxin 12/13 (Stx12) levels in the liver of female mice treated with rapamycin. In line with the findings in invertebrates, chronic rapamycin treatment significantly enhanced Stx12 protein levels in mouse liver (Fig. 6a), suggesting that the TORC1-associated regulation of Syntaxin 12/13 is conserved from yeast to mammals. To address whether reduced TORC1-S6K signalling also affects immunoageing in mice, we performed proteomics profiling of old liver samples from long-lived mice treated with rapamycin (Table 4) and compared these data to previously published transcriptome analyses of livers of aged mice with rapamycin treatment or S6K1 deficiency^11,49^ (Fig. 6b). Network propagation analysis^25^ of the 4340 proteins shared between the three datasets identified immune-related processes, including inflammation and leukocyte proliferation as commonly down-regulated by S6K deficiency or rapamycin treatment, while lipid-and translation-related processes were enhanced (Fig. 6c, Table 5). Taken together, these findings indicate that the effects of TORC1-S6K signalling on Syntaxin 12/13 expression and immune ageing are conserved between flies and mammals.

**Figure 6:**
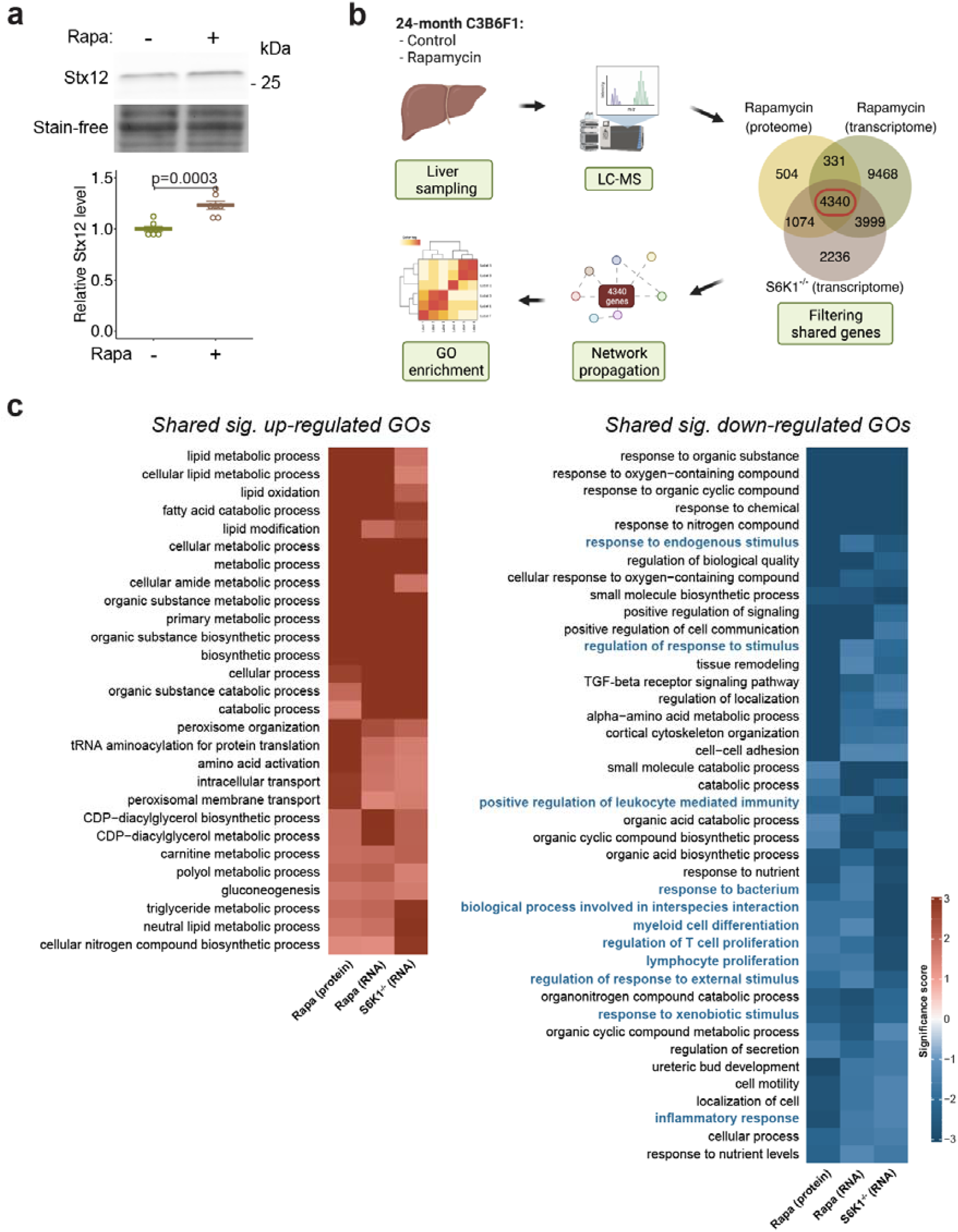
TORC1-S6K signalling regulates Syntaxin 12/13 expression and immune-related processes in mouse liver. **a**, Rapamycin treatment increased Syntaxin 12/13 (Stx12) level in the liver of 12-month-old mice (n=7, two-sided Student’s t-test). **b**, Schematic for the analysis workflow. Common genes present in all three datasets were used for network propagation and Gene Ontology (GO) analysis. **c**, Only GO terms significantly regulated (p<0.05) in the same direction in all three datasets are shown. Immune-related annotations are shown in blue. Cells are coloured by their log10 significance score.

## Discussion

The evolutionarily conserved mTOR signalling network plays an important role in determining longevity and is a potential target for geroprotective drugs in humans. S6K is a main downstream effector of mTOR signalling that has been implicated in regulation of lifespan in invertebrates and mammals. However, the molecular and cellular mechanisms underlying S6K-dependent longevity are still elusive. Here we showed that, in *Drosophila*, lowered activity of S6K in the fat body is essential for mTOR-dependent longevity. We further show that TORC1-S6K signalling regulates endolysosomal morphology, inflammageing and immunosenescence in the ageing fat body. Modifying endosome formation, but not autophagy, affected age-related inflammageing, suggesting a link between endosome formation and inflammageing. We identified the SNARE-like protein Syntaxin 13 (Syx13) as a molecular link that regulates both endosome formation and inflammageing downstream of TORC1-S6K signalling. Furthermore, we show that repression of the NF-κB like IMD pathway in the fly fat body enhances clearance of bacteria and extends lifespan. Importantly, we show that long-term treatment with rapamycin increases Syntaxin 12/13 (Stx12) levels in mouse liver and we identify alleviation of immune processes as a common denominator of TORC1-S6K inhibition in the liver of old mice, indicating that the effects of TORC1-S6K-Stx12 on immunoageing may be evolutionarily conserved from flies to mice. In summary, our findings highlight an important role for the TORC1-S6K-Syx13 signalling axis in inflammageing, immunosenescence and longevity (Fig. S1e).

Ubiquitous deletion of S6K increases lifespan in worms and of S6K1 does so in mice^11,23^. In *Drosophila*, overexpression of a kinase-dead version of S6K has also been shown to extend lifespan^50^. However, whether this lifespan extension was caused by a loss in S6K activity or by titration of the Raptor protein, a key component of the TORC1 complex that directly interacts with S6K via the Raptor-binding TOS motif, was not clear^51^. Here we show that RNAi-mediated downregulation of S6K is sufficient to increase lifespan in flies. Importantly, downregulation of S6K activity in the fat body but not in the intestine, brain, muscle, or heart extended lifespan to a similar extent as did ubiquitous downregulation, establishing the fat body as the key tissue for S6K mediated longevity in *Drosophila*. Consistent with this hypothesis, fat-body-specific overexpression of a constitutively active S6K protein blocked the effect of rapamycin on longevity, demonstrating that lowered S6K activity in the fat body is essential for TORC1-mediated lifespan extension. S6K and autophagy are both essential for lifespan extension upon TORC1 inhibition^4^. However, while S6K function is required in the fat body but not in the gut for TORC1 dependent longevity, autophagy is essential in the gut and not the fat tissue^8^. Thus, effector mechanisms downstream of TORC1 act in a tissue specific manner to regulate longevity, and the different, tissue-specific, health-enhancing effects of reduced TORC1 signalling collectively contribute to the powerful geroprotective effects of rapamycin.

Lysosomes are key to cellular proteostasis, and dysfunction of lysosomes contributes to age-associated pathologies^5^. Using lysotracker staining and electron microscopy we observed a strong increase in lysosomal size associated with multilamellar structures in fat bodies expressing activated S6K and treated with rapamycin. The morphological changes in the lysosome are reminiscent of lysosomal storage diseases (LSDs)^33^, and probably suggest impaired function of the lysosome caused by defects in lysosome or membrane fusion. This was further supported by the finding that TORC1-S6K dependent morphological changes in the lysosome were rescued by overexpression of Syntaxin 13 (Syx13), a SNARE family protein that regulates autophagosome maturation and vesicle fusion^52,53^. Knock down of Syx13 in human cells also induces multilamellar lysosome-like organelles^25^, suggesting that the function of Syx13 in regulating lysosomal morphology is evolutionarily conserved between flies and humans. Furthermore, Syx13 levels were upregulated in long-lived S6K mutant worms^23^ and also in various rapamycin-treated animals, including yeast^48^, flies and mice, suggesting that the regulation of Syx13 by TORC1 via S6K is evolutionarily conserved among these species. In contrast to our results from chronically rapamycin-treated mice, protein expression of Syx13 has been shown to be positively regulated by short-term TORC1 inhibition upon neuronal injury or in mammalian cell culture^54-56^, indicating a difference between acute stress and chronic inhibition of the mTOR pathway. The exact mechanism by which TORC1-S6K regulates Syx13 protein levels is currently unknown. Syx13 transcription was only mildly increased by rapamycin treatment in the fly fat body (Fig. S6b), suggesting that TORC1-S6K probably regulate Syx13 via a post-transcriptional mechanism.

Our data show that TORC1-S6K-Syx13 signalling regulates age-associated activity of the NF-κB like transcription factor Relish and AMP expression in the fly fat body, the main immune-responsive tissue of *Drosophila*. Importantly, S6K had no effect on Relish activity in young flies upon an acute bacterial challenge, suggesting that the effects of TORC1-S6K on the IMD pathway are indirect via regulation of other processes that change with age and then affect inflammation. Age-associated inflammation could be caused by external factors like pathogen stimulation due to increased gut leakiness with age and/or intrinsic deterioration of fat body function^44,57^. The finding that rapamycin repressed the number of Relish positive nuclei and AMP expression even when bacteria growth was prevented by antibiotic treatment suggests that TORC1-S6K-signalling regulates inflammageing via suppression of pathogen-independent sterile inflammation, which is in line with the findings in mammals^19,58^. Cellular senescence is a main driver of age-related sterile inflammation and can be caused by several factors^18^. Loss of nuclear Lamin has been associated with age-related senescence in the *Drosophila* fat body^37^. While we could confirm the age-related decline in Lamin levels in our proteomics dataset, neither rapamycin treatment nor S6K repression prevented the age-related loss of Lamin, suggesting that TORC1-S6K regulates age-related inflammation independent of Lamin. DNA damage is another driver of cellular senescence that increases with age in the fly fat body and amelioration of DNA damage specifically in the fat body increases lifespan in flies^59^. Furthermore, rapamycin treatment reduces senescence and improves immune function in DNA repair-deficient progeroid mice ^60^. Thus, reduced DNA damage by inhibition of mTOR in the fly fat body is a potential candidate mechanism that should be explored in the future to address how TORC1-S6K signalling regulates inflammageing. One of the most common triggers of inflammageing in mammals is the accumulation of adipose tissue during ageing^61^. Adipose tissue from old mice experiences endoplasmic reticulum (ER) stress and inflammation^62^; however, TORC1-S6K signalling did not affect the age-associated increase of ER stress indicator Hsc70-3, the ortholog of mammalian Grp78, in the fly fat body. Given that rapamycin suppresses ER-stress and its associated apoptosis in vitro^63^, the involvement of TORC1-S6K-Syx13 signalling in ER-stressed-dependent inflammageing needs further investigation.

Maintaining lysosomal function with age improves healthspan and lifespan in animals as diverse as yeast, worms, flies, and mice^6,8,64-66^. However, the effects have been mostly attributed to improvements in autophagy^5,67^ and not in endolysosomal function. Our results indicate that positive effects of reduced TORC1 signalling in the fly fat body do not rely on increased autophagy but on changes in the endolysosomal system. But how does the endolysosomal system affect age-related inflammation? One possibility is that the endolysosomal system is involved in the elimination of internal stimuli like cell debris and damage-associated molecular patterns (DAMPs), which accumulate during ageing and initiate pro-inflammatory immune signals^68,69^. Loss of phagocytosis or endocytosis capacity is a general age-associated alteration across various innate immune cells in mice and humans^70-72^, and may cause defects in tissue homeostasis, leading to chronic inflammation at advanced age. As Syx13 mediates endosome-lysosome fusion, overexpressing Syx13 may accelerate the internalisation and degradation of the internal stimulus and thereby suppress the age-associated immune hyperactivation. Alternatively, the endolysosomal system may directly affect the intracellular transport and degradation of immune regulators. Supporting this hypothesis, disrupting endosomal trafficking induces the accumulation of cytokine receptors and subsequently activates the IMD/NF-κB pathway in fly fat bodies^73^ and in human cells^74^. Thus, TORC1-S6K-Syx13 signalling may regulate inflammageing via regulating endosomal trafficking and the turnover of immune factors, a hypothesis that should be addressed experimentally in the future.

Another common feature of the aged immune system is immunosenescence^41,42^. Manipulation of TORC1-S6K-Syx13 signalling in the fly fat body alleviated pathogen clearance, suggesting suppression of immunosenescence in old flies. Immunosenescence has long been considered to drive inflammageing^60^. However, ameliorating inflammageing by repressing key inflammatory regulator Relish in the fly fat body is sufficient to ameliorate immunosenescence, suggesting that the inflammageing in the fly fat body acts upstream of immunosenescence. Immunosenescence in flies is often caused by the decreased phagocytic activity of haemocytes at advanced age^44,75^. Since the immune response in the fat body controls haemocyte activation^76^, one potential hypothesis would be that TORC1-S6K-Syx13 signalling in the fat body regulates haemocyte ageing by inter-tissue communication, thereby ameliorating immunosenescence. In addition to the haemocyte-related cellular immune response, the fly fat body secrets AMPs into the haemolymph to kill invading pathogens; however, the functional capacity of this humoral immune response declines with ageing^77^. Regulation of TORC1-S6K-Syx13 signalling in the fat body hence may be important for mediating pathogen-induced AMP expression in old flies.

In summary, our results establish that TORC1-S6K-Syx13 signalling regulates inflammageing in hepatic tissues via the endolysosomal system, thereby alleviating immunosenescence and enhancing longevity.

## Methods

### Fly stock and husbandry

All transgenic fly lines were backcrossed for at least six generations into the outbred wild-type strain, white Dahomey (*w*^*Dah*^)^78^. For experiments, flies were maintained on 10% (w/v) brewer’s yeast, 5% (w/v) sucrose and 1.5% (w/v) agar food at 25°C, 60% humidity, on a 12□h:12 h light:dark cycle. Rapamycin (LC Laboratories, #R-5000) was dissolved in ethanol and added to the food at a concentration of 200□µM. RU486 (Sigma, #M8046) was dissolved in ethanol and added to the food at a concentration of 100 µM to induce gene expression using the GeneSwitch system^79^. The corresponding control food contained only ethanol. Female flies were used in all experiments. Fly stocks are listed in Table 6. Fly genotypes of in each figure are indicated in the figure legend or body.

### Mouse husbandry

The mouse rapamycin study was performed in accordance with the recommendations and guidelines of the Federation of the European Laboratory Animal Science Association (FELASA), with all protocols approved by the Landesamt für Natur, Umwelt und Verbraucherschutz, Nordrhein-Westfalen, Germany (reference no. 81-02.04.2019.A313). Female F_1_ hybrid mice (C3B6F1) were generated in-house by crossing C3H/HeOuJ females with C57BL/6NCrl males (strain codes 626 and 027, respectively, Charles River Laboratories). Animals were housed in groups of five females in individually ventilated cages under specific-pathogen-free conditions with constant temperature (21□°C), 50–60% humidity and a 12□h:12 h light:dark cycle. For environmental enrichment, mice had constant access to nesting material and chew sticks. Rapamycin treatment was initiated at 6 months of age and was administrated continuously. Encapsulated rapamycin was obtained from Rapamycin Holdings and was added to the food (ssniff R/M-low phytoestrogen, ssniff Spezialdiäten) at a concentration of 42 mg/kg. Control food contained corresponding amounts of the encapsulation material Eudragit S100. Both rapamycin and control food contained 3.2 mL/kg PEG-400. For tissue collection, mice were killed by cervical dislocation, and tissues were rapidly collected and snap-frozen using liquid nitrogen.

### Generation of an *UAS-Syx13* transgenic fly line

To generate the transgenic *UAS-Syx13* fly line, full-length *Syx13* sequence was PCR amplified with primers (forward: *TTTTTTCTCGAGCACCatgtccaaggccttgaaca* and reverse: *TTTTTTTCTAGAttaactgttcagtttggcaacga*) using cDNA clone LD27581 (Drosophila Genomics Resource Center) as template. The PCR product was digested with *XhoI* and *XbaI* (NEB) and cloned into the *pUAST-attB* vector^80^. To increase expression efficiency, a *CACC* Kozak sequence was inserted directly before the *ATG* start codon. The phiC31-mediated integrase system^80^ was used to generate transgenic flies, using the *attP40* insertion site^81^.

### Lifespan, fecundity and starvation assays

For lifespan assays, parental flies were crossed in cages with grape juice agar plates and fresh yeast paste. Flies were allowed to lay eggs for 18 h, embryos were collected in PBS and dispensed into bottles at 20 µl per bottle to achieve standard larval density. Flies that enclosed within a 24 h time window were then transferred to fresh bottles where they were allowed to mate for 48 h. Subsequently, flies were anaesthetised with CO_2_ and 20 female flies were sorted to vials. Flies were transferred to fresh food vials every two to three days and scored for death. For fecundity assays, flies were treated as described above for the lifespan assay, but only five female flies were used per vials. Eggs laid within 20 h were collected and counted twice a week in the first four weeks. For starvation assays, female flies were reared and maintained as for lifespan assay. 10-day-old flies were transferred to 1% agar and scored for deaths at least twice per day.

### Peptide preparation for LC-MS/MS analysis

Dissected fat bodies from young (day 10) and old (day 50) female flies (five fat bodies were pooled as a replicate, four replicates per group) or 50 mg mouse livers were lysed in 6 M guanidinium chloride, 2.5 mM TCEP, 10 mM chloroacetamide, 100 mM Tris-HCl for 10 min at 95°C. Lysates were further disrupted by using a Bioruptor plus (Diagenode) with 30 s sonication and 30 s break for 10 cycles. Samples were then diluted 10fold with 20 mM Tris and trypsin (Trypsin Gold, Promega, #V5280, 1:200 w/w) and digested overnight at 37°C. Digestion was stopped by adding formic acid to a final concentration of 1%. Peptide cleaning was performed by using in-house C18-SD (Empore) StageTips^82^. Therefore, StageTips were washed with methanol and 40% acetonitrile (ACN)/0.1% formic acid (FA) and finally equilibrated with 0.1% FA. Digested peptides were loaded on the equilibrated StageTips. StageTips were washed twice with 0.1% FA and peptides were eluted with 40% ACN/0.1% FA and then dried in a Speed-Vac (Eppendorf) at 45°C for 45 min.

### TMT labelling and fractionation

Eluted peptides were reconstituted in 0.1M triethylammonium bicarbonate (TEAB). TMTpro 16plex (ThermoFisher, #A44522) labelling was carried out according to the manufacturer’s instruction with the following changes: 0.5 mg of TMT reagent was re-suspended with 33 µL of anhydrous ACN. 7 µL of TMT reagent in ACN was added to 9 µL of clean peptide in 0.1M TEAB. The final ACN concentration was 43.75% and the ratio of peptides to TMT reagent was 1:20. All 16 samples were labelled in one TMT batch. After 60 min of incubation the reaction was quenched with 2 µL of 5% hydroxylamine. Labelled peptides were pooled, dried, re-suspended in 0.1% FA, and desalted using StageTips. Samples were fractionated on a 1 mm x150 mm, 130Å, 1.7 µm ACQUITY UPLC Peptide CSH C18 Column (Waters, #186006935), using an Ultimate 3000 UHPLC (ThermoFisher). Peptides were separated at a flow of 30 µL/min with a 88 min segmented gradient from 1% to 50% buffer B for 85 min and from 50% to 95% buffer B for 3 min; buffer A was 5% ACN, 10 mM ammonium bicarbonate (ABC), buffer B was 80% ACN, 10 mM ABC. Fractions were collected every three minutes, and fractions were pooled in two passes (1 + 17, 2 + 18 … etc.) and dried in a Speed-Vac (Eppendorf).

### LC-MS/MS analysis and protein identification

Dried fractions were re-suspended in 0.1% FA and separated on a 500 mm, 0.075 mm Acclaim PepMap 100 C18 HPLC column (ThermoFisher, #164942) and analysed on a Orbitrap Lumos Tribrid mass spectrometer (ThermoFisher) equipped with a FAIMS device (ThermoFisher) that was operated in two compensation voltages, −50 V and −70 V. Synchronous precursor selection based MS3 was used for TMT reporter ion signal measurements. Peptides were separated by EASY-nLC1200 using a 90 min linear gradient from 6% to 31% buffer; buffer A was 0.1% FA, buffer B was 0.1% FA, 80% ACN. The analytical column was operated at 50°C.. Raw files corresponding to the mouse livers were split based on the FAIMS compensation voltage using FreeStyle (ThermoFisher). Fly fat body proteomics data were analysed using Proteome Discoverer (version 2.4.1.15); mouse liver proteomics data were analysed using MaxQuant (version 1.6.17.0). Isotope purity correction factors, provided by the manufacturer, were included in the analysis.

### Proteomics data analysis

Peptide intensity values were log2-and z-transformed. The results were rescaled by multiplying the z-intensity of each individual peptide with the global standard deviation of the log2-transformed data and adding back the global mean of the log2-transformed data. Only proteins that were detected in at least three out of the four replicates for each treatment were considered for downstream analyses. Missing values were imputed using the impute package (version 1.66.0) in R. Differential expression analysis was performed using the limma package (version 3.48.3) in R. For principal component analysis and plotting, batch effects from different dissection timepoints were removed from the normalized data using the limma package.

### Network propagation

High confidence interactions of the STRING protein-protein association network database for *Drosophila melanogaster* or *Mus musculus*^*83*^ were extracted as background for later propagation (combined score > 899). Network propagation was performed using the BioNetSmooth package (version 1.0.0) in R. Log2 fold changes of comparisons were imported into the network and propagated with α = 0.5 for 26 iterations (for flies) or 25 iterations (for mice). Top and bottom 5% proteins, which contain more than four protein-protein interactions were used for further analysis.

### Gene ontology term enrichment

Gene ontology information was retrieved from org.Dm.eg.db package (version 3.13.0) or Uniprot-GOA database (http://www.ebi.ac.uk/GOA/, version 2022-04-30) in R. The high confidence interactions of STRING database (combined score > 899) were set as background and the results of network propagation were used for gene ontology term enrichment analysis using Fisher tests with a minimal node size of five from ViSEAGO package (version 1.6.0) in R. For *Lsp2GS>S6K*^*CA*^ dataset, S6K^CA^-dependent Rapa-induced annotations were selected if the p-value of Rapa vs Control annotation is at least 100 times than the p-value of Induced *vs* Induced+Rapa annotation (EtOHvsRapa..log10_pvalue -RUvsRURapa..log10_pvalue > 2). S6K^CA^-independent Rapa-induced annotations were selected if the p-value of Rapa *vs* EtOH annotation is greater than 0.01 (EtOHvsRapa..log10_pvalue > 2) and if the p-value of Rapa *vs* EtOH annotation is not 100 times than the p-value of Induce *vs* Induced+Rapa annotation (EtOHvsRapa..log10_pvalue -RUvsRURapa..log10_pvalue ≤ 2). For mouse liver analysis, the biological processes significantly regulated (p<0.05) as the same direction in all three datasets (hepatic proteome of 24m rapamycin-treated mice, hepatic microarray of 25m rapamycin-treated mice^49^, and hepatic microarray of 20m S6K1 deficient mice^11^) are plotted as shared up-or down-regulated GO annotations after reducing redundancy via REVIGO (cutoff: “0.7”, valueType: “pvalue”, measure: “SIMREL”). The significance score was calculated by the -log10_p-value with the sign of regulation direction (positive values for up-regulated GOs and negative values for down-regulated GOs).

### Electron microscopy

Fat bodies were fixed in 4% Formaldehyde (Science Services) and 2,5% glutaraldehyde (Merck) in 0.1M Cacodylate buffer (AppliChem) for 48 h at 4°C. After washing in 0.1M cacodylate buffer, tissues were treated with 2% osmiumtetroxid (Science Services) in 0.1M Cacodylate buffer for 2 h. After dehydration of the sample with ascending ethanol concentrations followed by propylenoxid, samples were embedded in Epon (Sigma). Ultrathin sections (70 nm) were cut (EM-UC7, Leica Microsystems), collected onto mesh copper grids (Electron Microscopy Sciences), and contrasted with uranyl acetate (Plano GMBH) and lead citrate (Sigma). At least ten images per sample were acquired with a transmission electron microscope (JEM 2100 Plus, JEOL), a OneView 4K camera (Gatan) with DigitalMicrograph software at 80 KV at room temperature.

### Lysotracker staining

Tissues were dissected in PBS and stained with LysoTracker Red DND-99 (ThermoFisher, #L7528, 1:2000) and Hoechst 33342 (Enzo, #ENZ-52401, 0.5 ng/µl). Samples were rinsed with PBS and mounted with VECTASHIELD Antifade Mounting Medium (Vector Laboratories, #H-1000). Three images per sample were captured using a Leica TCS SP8 DLS confocal microscope with a 20x objective and 6x digital zoom in. Images were processed by background subtraction, median filtering, and spot quantification using Imaris 9 (Bitplane). Settings of the confocal microscope were kept consistent between images of an experiment. Lysostracker-positive puncta were defined as large, when the diameter was greater than 1.5 µm, which is the 75^th^ percentile of the diameter of rapamycin-induced Lysotracker-positive puncta.

### Immunofluorescence

Tissues were dissected in PBS and fixed 30 min with 4% formaldehyde, methanol-free (ThermoFisher, #28908). Samples were washed in 0.3% Triton-X/PBS (PBST), blocked in 5% bovine serum albumin (BSA)/PBST for 1 at room temperature, incubated in primary antibody overnight at 4°C, and in secondary antibody for 2 h at room temperature. The following primary antibodies were used: anti-Relish (Developmental Studies Hybridoma Bank, #21F3, 1:100), anti-Relish (RayBiotech, #RB-14-0004-200, 1:200). The following secondary antibodies were used: Alexa Flour 488 goat anti-mouse IgG (ThermoFisher, #A11001, 1:1,000), Alexa Flour 633 goat anti-mice IgG (ThermoFisher, #A21050, 1:1,000), Alexa Flour 594 goat anti-rabbit IgG (ThermoFisher, #A11012, 1:1,000), Alexa Flour 633 goat anti-rabbit IgG (ThermoFisher, #A21070, 1:1,000). Samples were mounted with VECTASHIELD Antifade Mounting Medium with DAPI (Vector Laboratories, #H-1200). Three images per sample were captured using a Leica TCS SP8 DLS confocal microscope with 20x objective and 6x digital zoom. For Relish protein localisation, images were processed by background subtraction and median filtering using Imaris 9 (Bitplane). The number of nuclei with high Relish fluorescence intensity was normalized by the total number of nuclei. Confocal settings were kept consistent between images of the same experiment.

### Immunoblotting

Tissues were homogenized and lysed in ice-cold RIPA buffer supplemented with cOmplete, Mini, EDTA-free, Protease Inhibitor Cocktail (Roche, #11836170001) and PhosSTOP phosphatase inhibitor tablet (Roche, #049068370001) using a hand-held homogenizer. Extracts were centrifuged and protein concentrations were determined using Pierce BCA Protein Assay Kit (ThermoFisher, #23225). Extracts were mixed with 4x Laemmli loading buffer and boiled for 5 min at 95°C. 10 µg (fly tissues) or 25 µg (mouse tissues) of proteins were loaded per lane on Any kD or 4-20% Criterion TGX stain-free precast gels (Bio-Rad) and transferred to Immobilon-FL PVDF membrane (Millipore, #IPFL00010). Membranes were blocked by Intercept TBS Blocking Buffer (LI-COR, #927-60001) for 1 h and probed with the following primary antibodies diluted in Intercept T20 TBS Antibody Diluent (LI-COR, #927-65001): anti-S6K^3^ (1:1,000), anti-pS6K T398 (Cell Signaling Technology, #9209, 1:1,000), anti-Relish (Developmental Studies Hybridoma Bank, #21F3, 1:100), anti-tubulin (Sigma, #T9026,1:5,000), anti-puromycin (Sigma, #MABE343, 1:1,000), anti-Stx12 (Synaptic Systems, #110 132, 1:1000). The following secondary antibodies were used: IRDye 800CW Goat anti-Mouse IgG (H + L) (LI-COR, #926-32210, 1:15,000) and IRDye 680RD Goat anti-Rabbit IgG (H + L) (LI-COR, #926-68071, 1:15,000). Total protein on the membrane was visualized as stain-free signal using ChemiDoc MP Imagers (Bio-Rad). Immunoblotting images were captured using Odyssey Infrared Imaging system with application software V3.0.30 (LI-COR) and were analysed using Fiji^84^ (US National Institutes of Health).

### RNA isolation and qRT-PCR

Total RNA was extracted from tissues using the RNeasy Mini Kit (Qiagen, #74106) following the manual. RNA concentration was measured using the Qubit RNA Broad Range Assay Kit (ThermoFisher, #Q10211). cDNA synthesis was performed using SuperScript VILO Master Mix (ThermoFisher, #11755-250) with 600 ng total RNA as input. qRT-PCR was performed using SYBR Green Master Mix (ThermoFisher, #4367659) on a QuantStudio6 Flex Real-Time PCR System (ThermoFisher).

Relative expression was determined using the ΔΔCT method and *Rpl32* as normalization control. The following primers were used:

*DptA-forward: CCACGAGATTGGACTGAATG*

*DptA-reverse: GGTGTAGGTGCTTCCCACTT*

*Syx13-forward: GCGGCAGGTCGAGCAAATA*

*Syx13-reverse: AGTTCCGAGGTGCCATCCT*

*Rpl32-forward: ATATGCTAAGCTGTCGCACAAATGG*

*Rpl32-reverse: GATCCGTAACCGATGTTGGGCA*

### Triacylglyceride (TAG) assay

Five whole flies were homogenized by 0.05% Tween-20 and incubated at 70°C for 5 min. Extracts were centrifuged, the supernatant was mixed with prewarmed Infinity Triglycerides reagent (ThermoFisher, #TR22421) and incubated at 37°C for 5 min. Protein content was determined by Pierce BCA Protein Assay Kit (ThermoFisher, #23225). Triglyceride levels were determined by plate reader at the absorbance of 540 nm using a triglyceride standard (Sigma, #17811-1AMP). Protein levels were used for normalization.

### Puromycin incorporation assay

Fat bodies were dissected in PBS and immediately transferred to Schneider’s Drosophila media (Biowest, #L0207-500) with 5 µg/ml puromycin (Sigma, #P8833). For negative control, no puromycin was added in the Schneider’s Drosophila media. Sample were incubated for 45 min with 400 rpm at 25°C and then lysed according to the immunoblotting assay.

### Antibiotic treatment

For antibiotic treatment, tetracycline and ampicillin were dissolved in water and added to the food in a concentration of 100 mg/l and 50 mg/l, respectively. In order to test whether the treatment was efficient at removing bacteria, flies were dipped in 70% ethanol for 3 min and rinsed with sterile PBS. Individual flies were mashed in sterile PBS. Samples were plated on LB plates, cultured at room temperature, and the number of bacterial colonies was scored at day 7.

### Infection assays

For infection assays, flies were infected by pricking the thorax with a fine needle dipped in freshly cultured *Ecc15* (OD^600^ = 200). At 0 or 12 hour post-infection (h.p.i.), flies were surface sterilized with 70% ethanol, rinsed with sterile PBS, and then three flies were homogenized in sterile PBS by FastPrep-24 Matrix-D. Tenfold-series dilutions of fly homogenates were plated on LB plates as spots, cultured at room temperature, and the number of bacterial colonies was scored the next day. 0 h.p.i. represented the initial infectious dose and was not included in the statistical analysis. For *ex vivo* infection assay, fat bodies were dissected in sterile PBS and immediately incubated with PBS containing freshly cultured *Ecc15* (final concentration OD^600^ = 0.2) at room temperature for 1 h. For sham control, tissues were incubated with sterile PBS at room temperature for 1 h. Samples were then processed as described for immunofluorescence assay.

### Statistics

All statistical analyses were performed in GraphPad Prism 9 and R 4.1.0. Linear mixed model and negative binomial generalized linear model were generated and analysed in R using lme4, lmertest, and emmeans package. Proteomics data were analysed in R using impute, limma, ViSEAGO package. Cox Proportional Hazards (CPH) test were performed in R using survival package. Log-rank tests were performed in Microsoft Excel for Mac (version 16.62) and R using survival package. For small sample sizes (n < 8), data are presented as individual points with mean□±□s.e.m.. For large sample size (n ≥ 8), box plots were used with median, 25^th^ and 75^th^ percentiles, and Tukey whiskers indicated. Sample sizes and statistical tests used are indicated in the figure legend.

## Acknowledgements

We thank Gábor Juhász for the Lamp1 reporter fly strains and Jacqueline Eßer for generating transgenic flies. Christian Kukat and the FACS and Imaging Core Facility at the Max Planck Institute for Biology of Ageing are acknowledged for their support with confocal microscopy. Astrid Schauss, Felix Gaedke, and Janine Heise from the Imaging Core Facility at CECAD (Cologne) are acknowledged for generating electron microscopy data. We acknowledge Xinping Li, Thomas Colby and Ilian Atanassov from the Proteomics Core Facility at the Max Planck Institute for Biology of Ageing for generating the proteomics data. We thank the Bloomington Drosophila Stock Center, the Vienna Drosophila Resource Center (VDRC), FlyORF, the Drosophila Genomics Resource Center (DGRC), and the Developmental Studies Hybridoma Bank (DSHB) for fly strains and reagents. This project has received funding from the European Research Council (ERC) under the European Union’s Horizon 2020 research and innovation programme (grant agreement no. 741989) and the Max-Planck-Gesellschaft. Pingze Zhang was supported by a fellowship from the Chinese Scholarship Council. All schematic diagrams were created with BioRender.com.

## Author contributions

P.Z., S.G. and L.P. conceived and designed the study. P.Z. and J.C. conducted experiments. P.Z. analysed the data. P.Z., S.G. and L.P. wrote the manuscript.

## Competing interests

Authors declare no competing interests.

## Corresponding authors

Correspondence to Sebastian Grönke and Linda Partridge.

## Notes

### Competing Interest Statement

The authors have declared no competing interest.

