## Supplementary material for "Inhibition of S6K lowers age-related inflammation and immunosenescence and increases lifespan through the endolysosomal system": Fig. S1

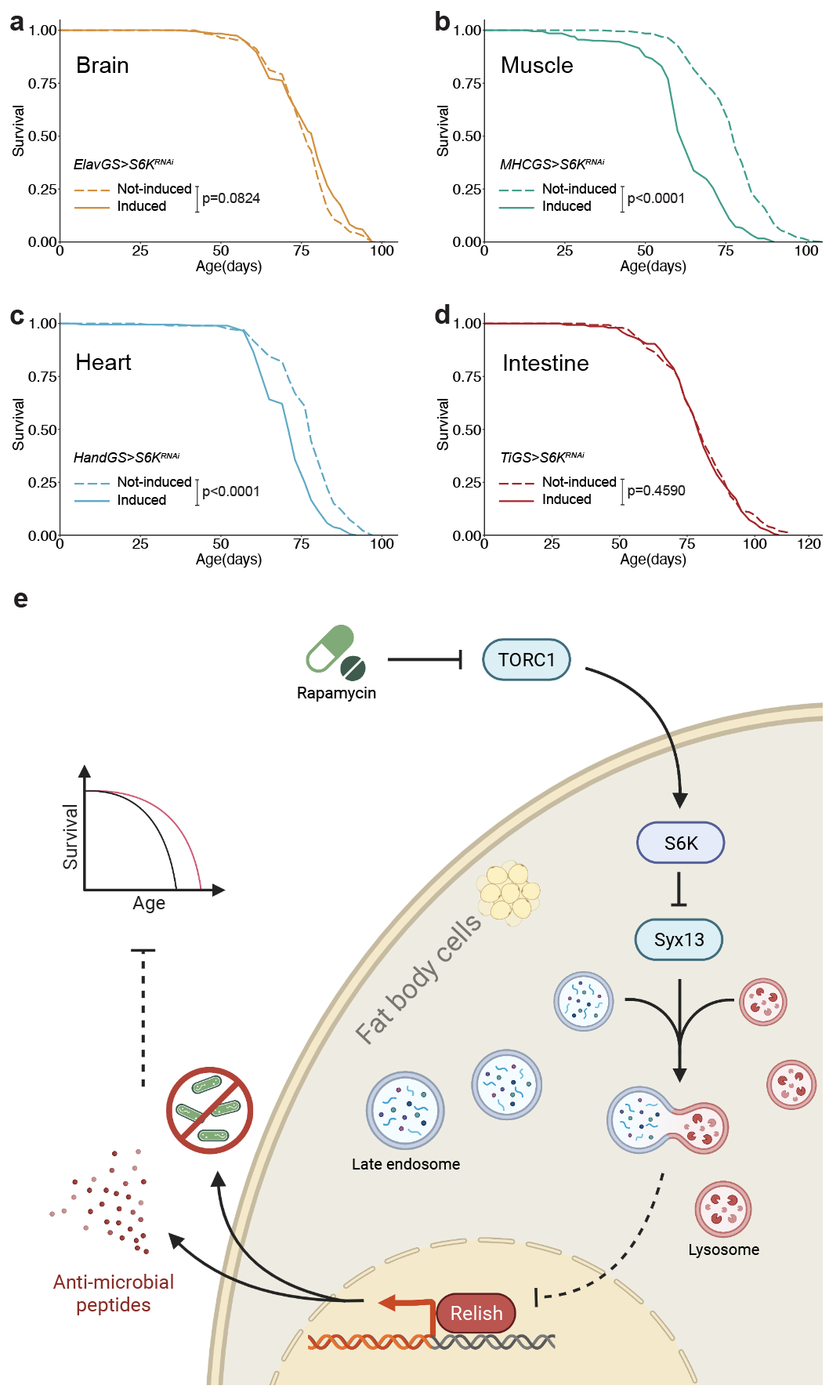

**Figure S1: Downregulation of S6K activity in the brain, muscle, heart, or intestine does not extend lifespan and the schematic of the findings. a,** Adult-onset repression of S6K in the brain using *elavGS>S6K^RNAi^* did not affect lifespan (n=200). **b,** Adult-onset repression of S6K in the muscle using *MHCGS>S6K^RNAi^* shortened lifespan (n=200). **c,** Adult-onset repression of S6K in the heart using *HandGS>S6K^RNAi^* shortened lifespan (n=200). **d,** Adult-onset repression of S6K in the intestine using *TiGS>S6K^RNAi^* did not affect lifespan (n=150). Log-rank test. **e,** Schematic of the findings.

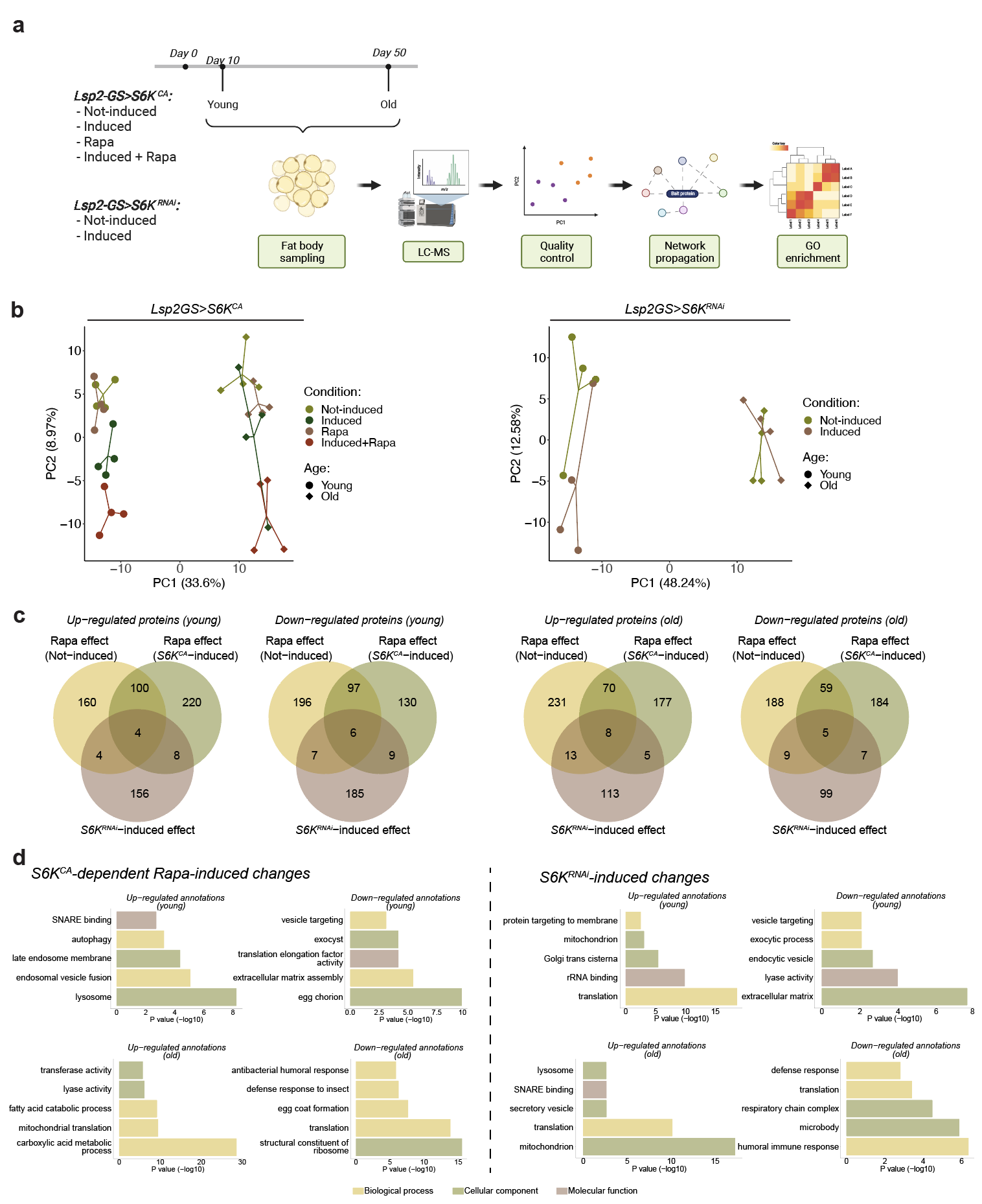

**Figure S2: Proteomics design and analysis. a,** Schematic for 16-plex TMT based proteomics experimental design. **b,** Principal component analysis projections of all proteomic replicates of *Lsp2GS>S6K^CA^* (left) and *Lsp2GS>S6K^RNAi^* (right) dataset, showing clear separation of the conditions and the ages. Each proteomics replicate represents a pool of five fat bodies**. c,** Venn diagram showing the number of up- and down- regulated proteins (p<0.05) of each comparison at each time point. Not-induced Rapa effect represented the differentially expressed proteins in the comparison of *S6K^CA^ not-induced vs S6K^CA^ not-induced+Rapa*; *S6K^CA^-*induced Rapa effect represented the differentially expressed proteins in the comparison of *S6K^CA^-induced vs S6K^CA^-induced+Rapa;* and *S6K^RNAi^-*induced effect represented the differentially expressed proteins in the comparison of *S6K^RNAi^ not-induced vs S6K^RNAi^-induced.* **e,** Representation of Gene Ontology (GO) terms in *Lsp2GS>S6K^CA^* dataset (left) and *Lsp2GS>S6K^RNAi^*  dataset (right) after network propagation. S6K^CA^-dependent Rapa-induced annotations were selected if the p-value of *S6K^CA^ not-induced* *vs S6K^CA^ not-induced*+*Rapa* annotation is at least 100 times greater than the p-value of *S6K^CA^-induced* *vs S6K^CA^-induced+Rapa* annotation (EtOHvsRapa..log10_pvalue - RUvsRURapa..log10_pvalue > 2). Colour indicates the categories of the terms.

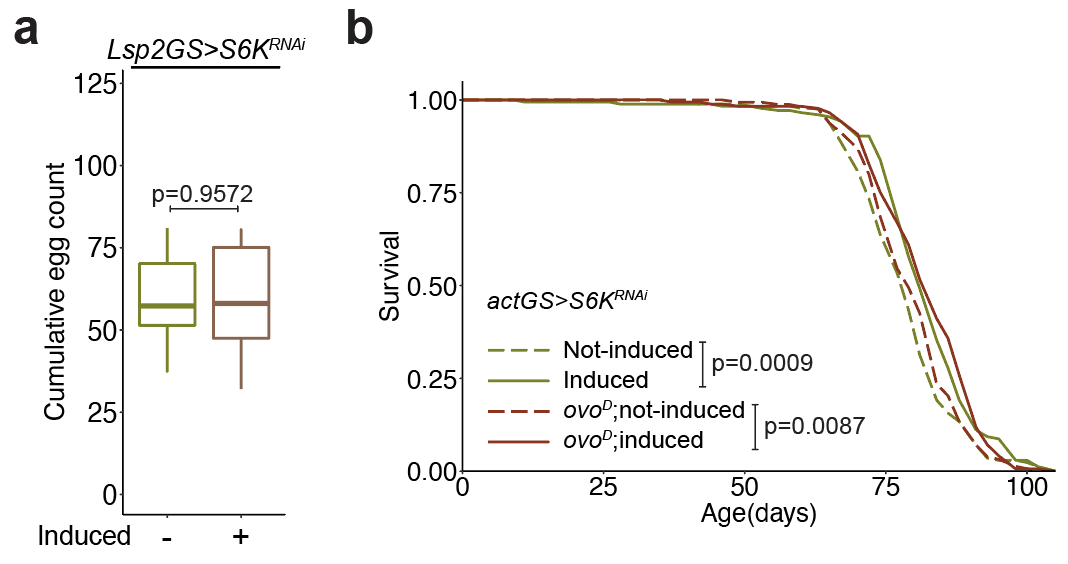

**Figure S3: S6K-related longevity is independent of reproduction. a,** Adult-onset repression of S6K in the fat body using *Lsp2GS*>S6K^RNAi^ did not affect cumulative egg laying of flies from day 3 to day 25 (n=10). **b,** RNAi-mediated knockdown of S6K expression using *actGS>S6K^RNAi^* extended lifespan in both control background and sterile *ovo^D^* background (*ovo^D^*: p=0.5371, *actGS>S6K^RNAi^* induction: p=0.0015, interaction p=0.6719, n=180). Data are displayed as Tukey box plot **(a).** Each data point represents an average value per five flies **(a).** Two-sided Student’s t-test **(a)** or log-rank test and CPH test **(b)**.

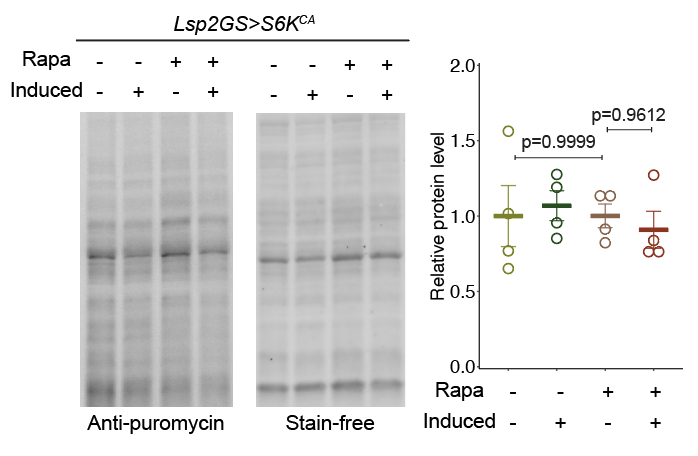

**Figure S4: Rapamycin treatment or overexpression of constitutively active S6K does not regulate** **global translation activity in the fat body.** Rapamycin treatment or overexpression of constitutively active S6K specifically in the fat body (*Lsp2GS>S6K^CA^*) did not affect newly synthesized protein in the fat body of young flies, depicted by puromycin immunoblotting (rapamycin: p=0.5660, *Lsp2GS>S6K^CA^* induction: p=0.9320, interaction p= 0.5610, n=4). Stain-free blot served as loading control. Data are mean ± s.e.m.. Each data point represents an average value per five fat bodies. Linear mixed model followed by Tukey’s multiple comparison test.

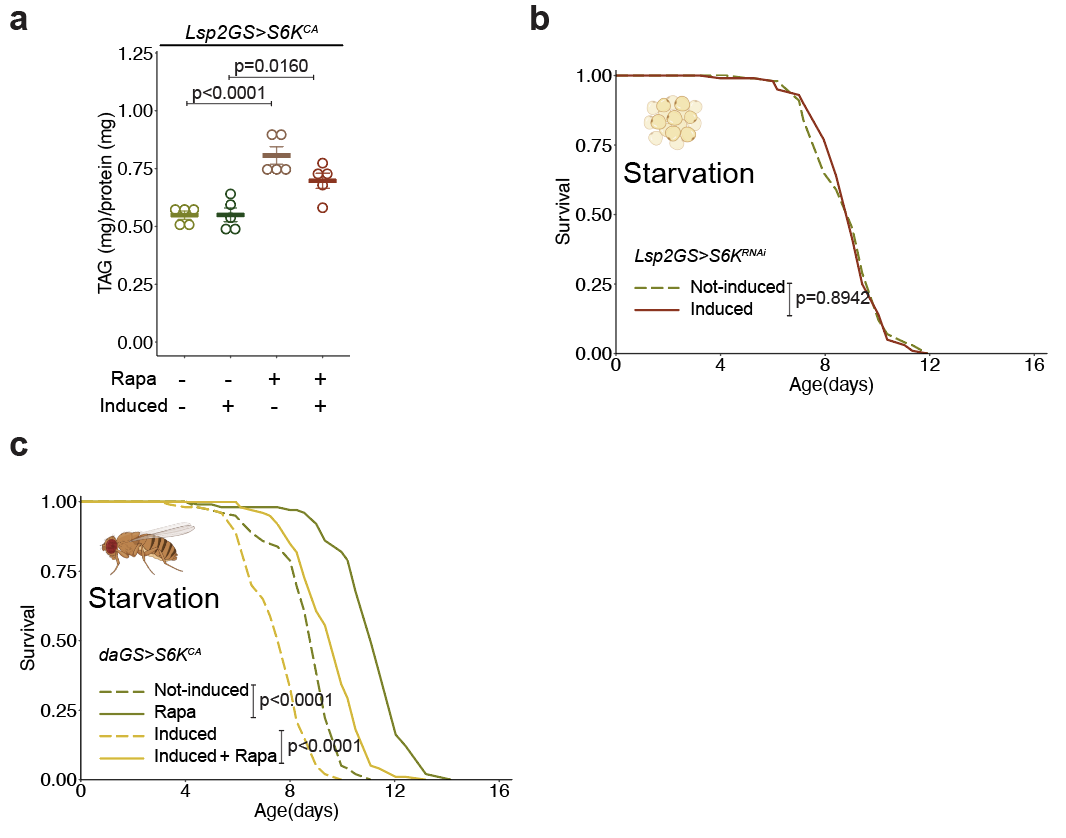

**Figure S5: Overexpression of constitutively active S6K does not block the effect of rapamycin on lipid homeostasis in the fly fat body. a,** Rapamycin increased triglyceride (TAG) storage of young flies. Expression of constitutively active S6K in the fat body (*Lsp2GS>S6K^CA^*) did not block the effect of rapamycin on triglyceride storage (rapamycin: p<0.0001, *Lsp2GS>S6K^CA^* induction: p=0.0957, interaction p=0.0890, n=5). **b,** Starvation resistance was not affected by RNAi mediated downregulation of S6K activity in the fat body (*Lsp2GS>S6K^RNAi^*, n=100). **c,** Rapamycin increased starvation resistance of young flies. Expression of constitutively active S6K ubiquitously (*daGS>S6K^CA^*) did not block the effect of rapamycin on starvation resistance (rapamycin: p<0.0001, *daGS>S6K^CA^* induction: p<0.0001, interaction p=0.5770, n=100). Data are mean ± s.e.m. **(a)**. Each data point represents an average value per five whole flies **(a)**. Linear mixed model followed by Tukey’s multiple comparison test **(a)**; log-rank test and CPH analysis **(b-c)**.

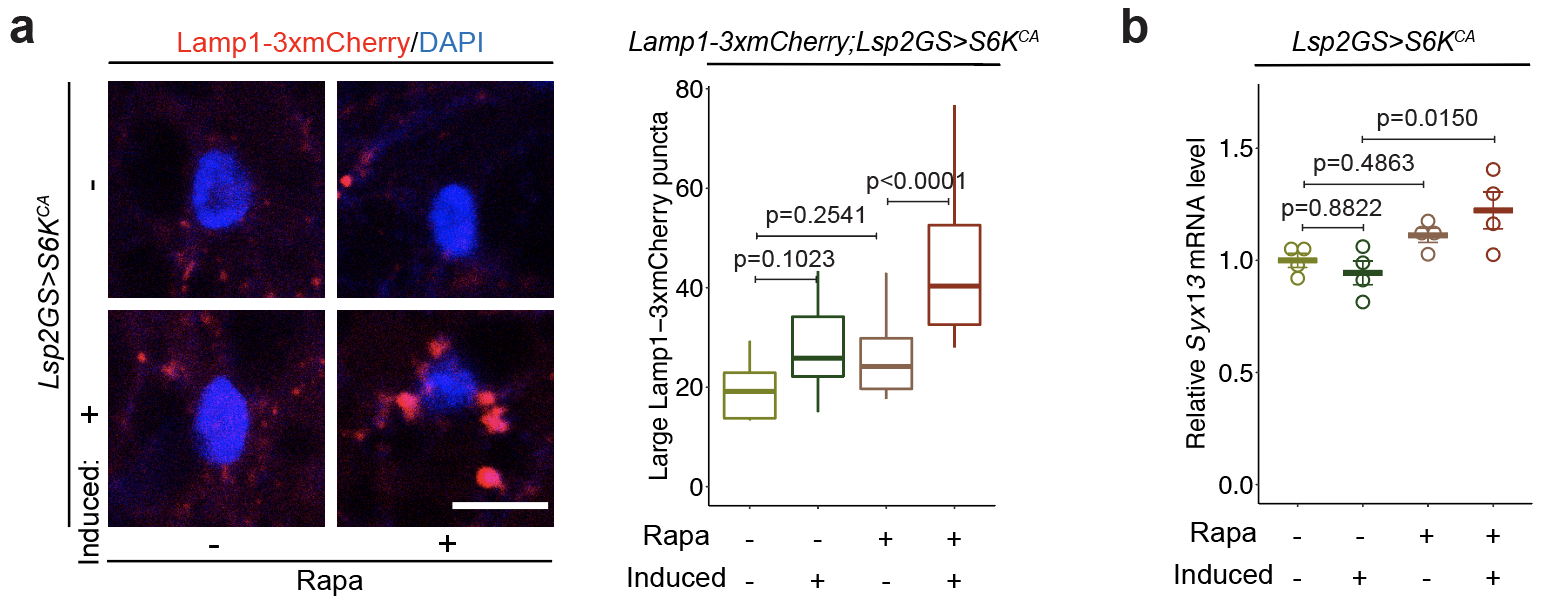

**Figure S6: TORC1-S6K signalling affects lysosomal size and Syx13 transcription in the fly fat body.** **a,** Visualization of enlarged lysosomes in the fat body of young flies treated with rapamycin and overexpressing constitutively active S6K (*Lsp2GS>S6K^CA^*) using a *3xmCherry-Lamp1* reporter (rapamycin: p<0.0001, *Lsp2GS>S6K^CA^* induction: p<0.0001, interaction p=0.0468, n=14). **b,** Fat body-specific adult-onset overexpression of constitutively active S6K significantly increased Syx13 transcription in response to rapamycin treatment in young fat bodies (rapamycin: p=0.1467, *Lsp2GS>S6K^CA^* induction: p=0.6155, interaction p=0.0035, n=5). Data are displayed as Tukey box plot **(a)** or mean ± s.e.m. **(b)**. Each data point represents an average value per fat body **(a)** or per five fat bodies **(b)**. Scale bar, 10 μm. Linear mixed model **(a)** or two-way ANOVA **(b)** followed by Tukey’s multiple comparison test.

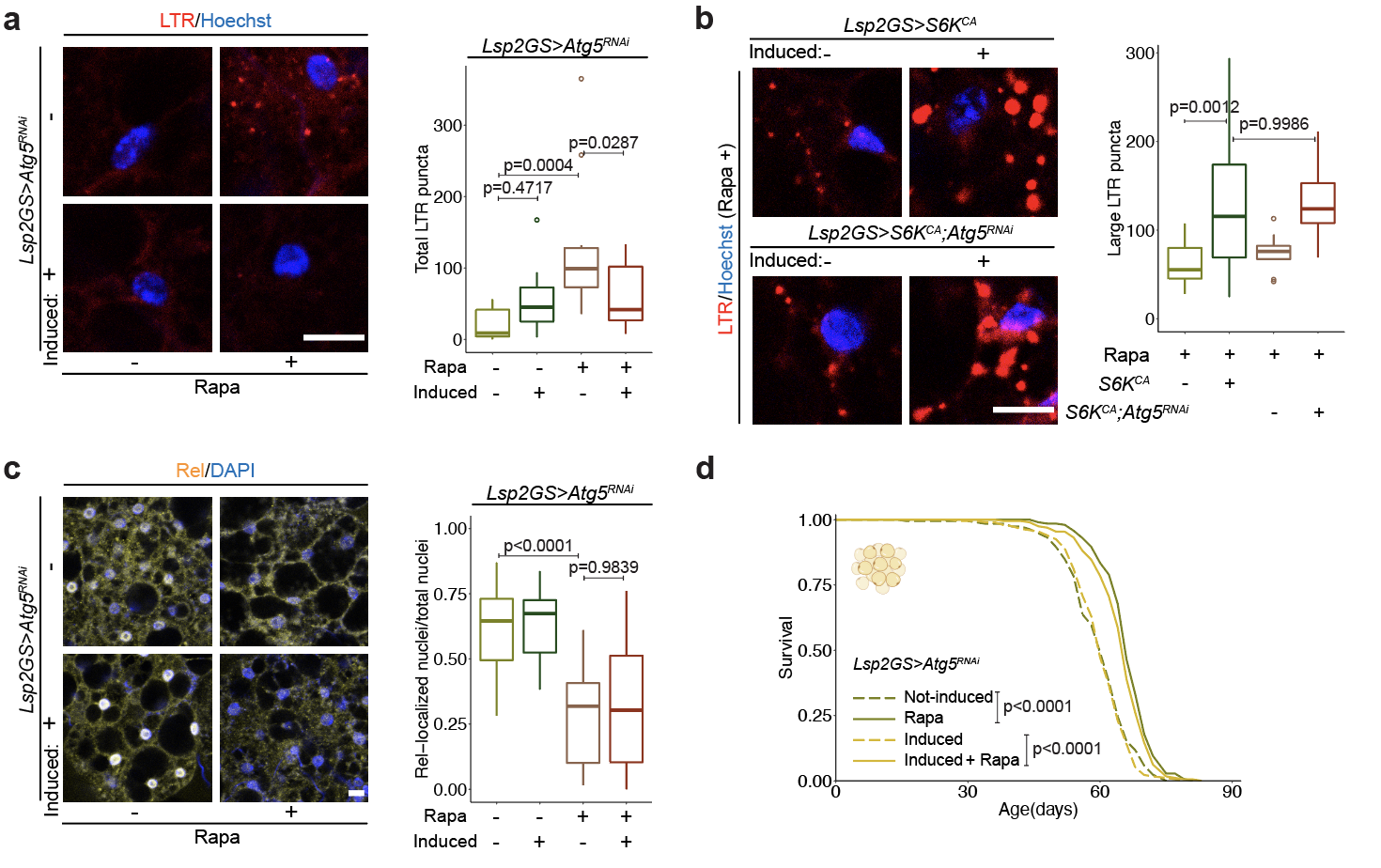

**Figure S7: Autophagy does not regulate lysosomal morphology or rapamycin-related health benefits. a,** Rapamycin treatment induced acid organelles in the fat body of young flies. RNAi mediated knock down of Atg5 specifically in the fat body (*Lsp2GS>Atg5^RNAi^*) blocked this phenotype, depicted by lysotracker staining (rapamycin: p=0.0025, *Lsp2GS>Atg5^RNAi^* induction: p=0.2947, interaction p=0.0035, n=12). **b,** RNAi mediated knock down of Atg5 specifically in the fat body (*Lsp2GS>S6K^CA^;Atg5^RNAi^*) did not affect the enlarged lysosome phenotype of fat body tissue of S6K overexpression flies (*Lsp2GS>S6K^CA^*) treated with rapamycin, depicted by lysotracker staining (*Lsp2GS>Atg5^RNAi^* induction: p=0.6766, *Lsp2GS>S6K^CA^* induction: p<0.0001, interaction p=0.5220, n=12). **c,** RNAi mediated knock down of Atg5 specifically in the fat body (*Lsp2GS>Atg5^RNAi^*) did not affect the effect of rapamycin on Relish localisation in the old fat body (rapamycin: p<0.0001, *Lsp2GS>Atg5^RNAi^* induction: p=0.6189, interaction p=0.9932, n=14). **d,** RNAi mediated knock down of Atg5 specifically in the fat body (*Lsp2GS>Atg5^RNAi^*) did not affect the effect of rapamycin on lifespan (rapamycin: p<0.0001, *Lsp2GS>Atg5^RNAi^* induction: p=0.4130, interaction p=0.4320, n=200). Data are displayed as Tukey box plot. Each data point represents an average value per fat body. Scale bar, 10 μm. Linear mixed model followed by Tukey’s multiple comparison test **(a-c)** or log-rank test and CPH analysis **(d)**.

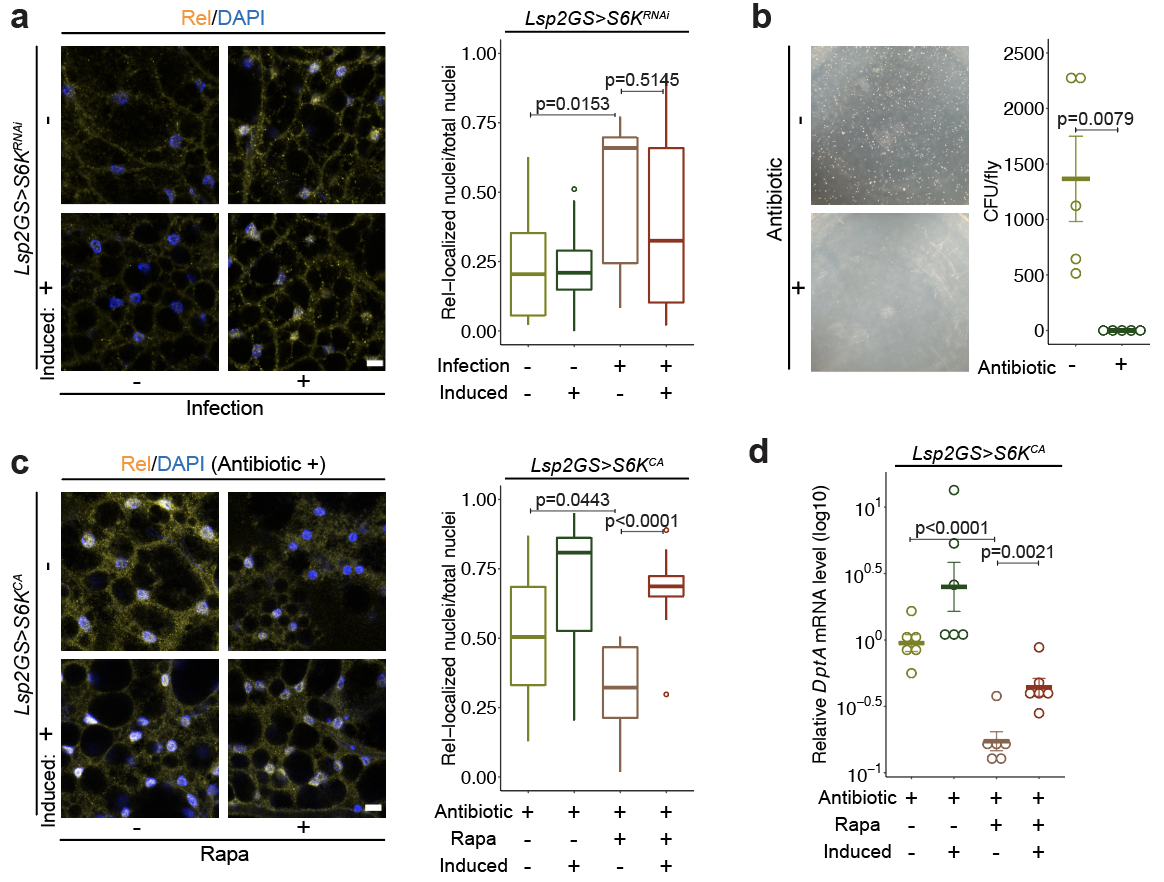

**Figure S8: TORC1-S6K-related benefits on inflammageing is independent on bacterial infection.** **a,** Bacteria infection by *Ecc15* induced accumulation of Relish in the nucleus in the young fat body. RNAi-mediated knockdown of S6K (*Lsp2GS>S6K^RNAi^*) did not affect the infection-induced Relish localisation (infection: p=0.0013, *Lsp2GS>S6K^RNAi^* induction: p=0.3499, interaction p=0.3151, n=12). **b,** Colony forming unit (CFU) assay shows the effectiveness of the antibiotic treatment using tetracycline and ampicillin to suppress bacterial growth in the treated flies (n=5). **c,** Rapamycin treatment suppressed age-related accumulation of Relish in the nucleus in the fat body of old flies treated with antibiotic. Overexpression of constitutively active S6K (*Lsp2GS>S6K^CA^*) blocked the effect of rapamycin on Relish localisation (rapamycin: p=0.0264, *Lsp2GS>S6K^CA^* induction: p<0.0001, interaction p=0.1300, n=14). **d,** Rapamycin treatment suppressed *DptA* expression in old fat body cells. Overexpression of constitutively active S6K (*Lsp2GS>S6K^CA^*) induced *DptA* expression of files treated with rapamycin and antibiotic (n=6). Data are displayed as Tukey box plot **(a, c)** or mean ± s.e.m. **(b, d)**. Each data point represents an average value per fat body **(a, c),** per whole fly **(b),** or per five fat bodies **(d**)**.** Scale bar, 10 μm. Linear mixed model followed by Tukey’s multiple comparison test **(a, c)**; Mann-Whitney test **(b)**; or two-sided Student’s t-test with log transformation **(d)**.

**Table 1**

List of proteins identified from TMT-based proteomic analysis in young (day 10) and old (day 50) fat bodies of *Lsp2GS>S6K^CA^* or *Lsp2GS>S6K^RNAi^* flies.

**Table 2**

List of proteins that differentially regulated by TORC1-S6K signalling. The proteins that were significantly changed (p<0.05) by both S6K knockdown and rapamycin treatment but were not altered by rapamycin treatment under S6K activation condition (p≥0.05) were selected as TORC1-S6K-dependent proteins.

**Table 3**

List of rapamycin-induced, S6K-dependent GO terms and S6K inhibition-induced GO terms. Results contain three sub-ontologies: biological process (BP), cellular component (CC) and molecular function (MF).

**Table 4**

List of proteins identified from TMT-based proteomic analysis in the liver of 24-month-old rapamycin-treated mice.

**Table 5**

List of GO terms aggregated across datasets (rapamycin-proteome, rapamycin-transcriptome, and S6K1^-/-^-transcriptome). Common genes which have been detected in all three datasets were used for network propagation and the following Gene Ontology (GO) analysis. Only functions significantly regulated (p<0.05) as the same direction in all three datasets are shown.

**Table 6**

List of fly strains used in this study.
